# Single cell resolution regulatory landscape of the mouse kidney highlights cellular differentiation programs and renal disease targets

**DOI:** 10.1101/2020.05.24.113910

**Authors:** Zhen Miao, Michael S. Balzer, Ziyuan Ma, Hongbo Liu, Junnan Wu, Rojesh Shrestha, Tamas Aranyi, Amy Kwan, Ayano Kondo, Marco Pontoglio, Junhyong Kim, Mingyao Li, Klaus H. Kaestner, Katalin Susztak

## Abstract

Determining the epigenetic program that generates unique cell types in the kidney is critical for understanding cell-type heterogeneity during tissue homeostasis and injury response.

Here, we profiled open chromatin and gene expression in developing and adult mouse kidneys at single cell resolution. We show critical reliance of gene expression on distal regulatory elements (enhancers). We define key cell type-specific transcription factors and major gene-regulatory circuits for kidney cells. Dynamic chromatin and expression changes during nephron progenitor differentiation demonstrated that podocyte commitment occurs early and is associated with sustained *Foxl1* expression. Renal tubule cells followed a more complex differentiation, where *Hfn4a* was associated with proximal and *Tfap2b* with distal fate. Mapping single nucleotide variants associated with human kidney disease identified critical cell types, developmental stages, genes, and regulatory mechanisms.

We provide a global single cell resolution view of chromatin accessibility of kidney development. The dataset is available via interactive public websites.

## Introduction

The mammalian kidney maintains fluid, electrolyte, and metabolite balance of the body and plays an essential role in blood pressure regulation and red blood cell homeostasis. The human kidney makes roughly 180 liters of primary filtrate each day that is then reabsorbed and modified by a long tubule segment. To perform this highly choreographed and sophisticated function, the kidney contains close to 20 highly specialized epithelial cells. The renal glomerulus acts as a 60 kD size-selective filter. The proximal part of the tubules is responsible for reclaiming more than 70% of the primary filtrate, which is done via unregulated active and passive paracellular transport ^1^, while the loop of Henle plays an important role in concentrating the urine. The distal convoluted tubule is critical for regulated electrogenic sodium reabsorption and potassium secretion. The last segment of kidney tubules is the collecting duct where the final concentration of the urine is determined via regulation of water channels as well as acid or base secretion. Understanding the development of these diverse cell types in the kidney is essential to understand kidney homeostasis, disease, and regeneration.

The mammalian kidney develops from the intermediate mesoderm via a complex interaction between the ureteric bud and the metanephric mesenchyme ^2^. In the mouse kidney, Six2 marks the self-renewing nephron progenitor population ^3^. The nephron progenitors commit and undergo a mesenchymal-to-epithelial transformation giving rise to the renal vesicle ^3^. The renal vesicle then undergoes segmentation and elongation, giving rise to epithelia from the podocytes to the distal convoluted tubules, while the ureteric bud becomes the collecting duct. Unbiased and hypothesis-driven studies have highlighted critical stages and drivers of early kidney development ^4^, that have been essential for the development of *in vitro* kidney organoid differentiation protocols ^5–7^. However, cells in organoids are still poorly differentiated, improving cellular differentiation and maturation of these structures remains a major challenge ^8^. Thus, the understanding of late kidney development, especially the cell type-specific driver transcription factors (TFs) is of great importance ^9–11^. Alteration in Wnt, Notch, Bmp, and Egf signaling significantly impacts cellular differentiation, but only a handful of TFs that directly drive the differentiation of distinct segments have been identified, such as *Pou3f3*, *Lhx1*, *Irx2*, *Foxc2,* and *Mafb* ^12^. Further understanding of the terminal differentiation program could aid the understanding of kidney disease development.

While single cell RNA sequencing (scRNA-seq) has improved our understanding of kidney development in mice and humans ^9,10,13,14^, it provides limited information of TFs, which are usually lowly expressed. Equally difficult is to understand how genes are regulated from scRNA-seq data alone. Chromatin state profiles, on the other hand, provide valuable insight to gene regulation mechanisms during cell differentiation, since they show not only the accessibility of the gene transcription start site (TSS), but also of distal regulatory regions such as enhancers. It is believed that enhancers are critical for establishing the cell type-specific gene expression pattern, but it has not been shown conclusively on a single cell level. Together with gene expression, open chromatin profiles can define the gene regulatory logic, which is the fundamental element of cell identity. However, there is a scarcity of open chromatin information by Assay for Transposase-Accessible Chromatin using sequencing (ATAC-seq) or chromatin immunoprecipitation (ChIP) data by ChIP-seq related to kidney development. In addition, epigenetic changes observed in bulk analyses mostly represent changes in cell composition, rather than cell type-specific changes ^15^, making it challenging to interpret bulk ATAC-seq data.

To this end, here we generated a single cell open chromatin and corresponding expression survey for the developing and adult mouse kidney, which will be available for the community via a searchable website (susztaklab.com/developing_adult_kidney/snATAC/ for snATAC-seq data, susztaklab.com/developing_adult_kidney/scRNA/ for scRNA-seq data, and susztaklab.com/developing_adult_kidney/igv/ for IGV view of peak tracks). Using this atlas, we have produced a new epigenome-based classification of developing and mature cells and defined cell type-specific regulatory networks. We also investigated key TFs and cell-cell interactions associated with developmental cellular transitions. Finally, we used the single cell open chromatin information to pinpoint putative target genes and cell types of several chronic kidney disease noncoding genome-wide association study (GWAS) loci.

## Results

### Single cell accessible chromatin landscape of the developing and adult mouse kidneys

To characterize the accessible chromatin landscape of the developing and adult mouse kidneys at single cell resolution, we performed single nuclei ATAC-seq (snATAC-seq) on kidneys of mice on postnatal day 0 (P0) at 3 and 8 weeks of age (**Figure 1a, Methods**). In parallel, we also performed bulk (whole kidney) ATAC-seq analysis at matched developmental stages. Following sequencing, we aggregated all high-quality mapped reads in each sample irrespective of barcode. The combined snATAC-seq dataset from all samples showed the expected insert size periodicity (**Figure S1a**) with a strong enrichment of signal at Transcription Start Sites (TSS, **Figure S1b**), indicating high data quality. The snATAC-seq data showed high concordance with the bulk ATAC data (Spearman correlation coefficient >0.84, **Methods**, **Figure S1c**).

**Figure 1.**
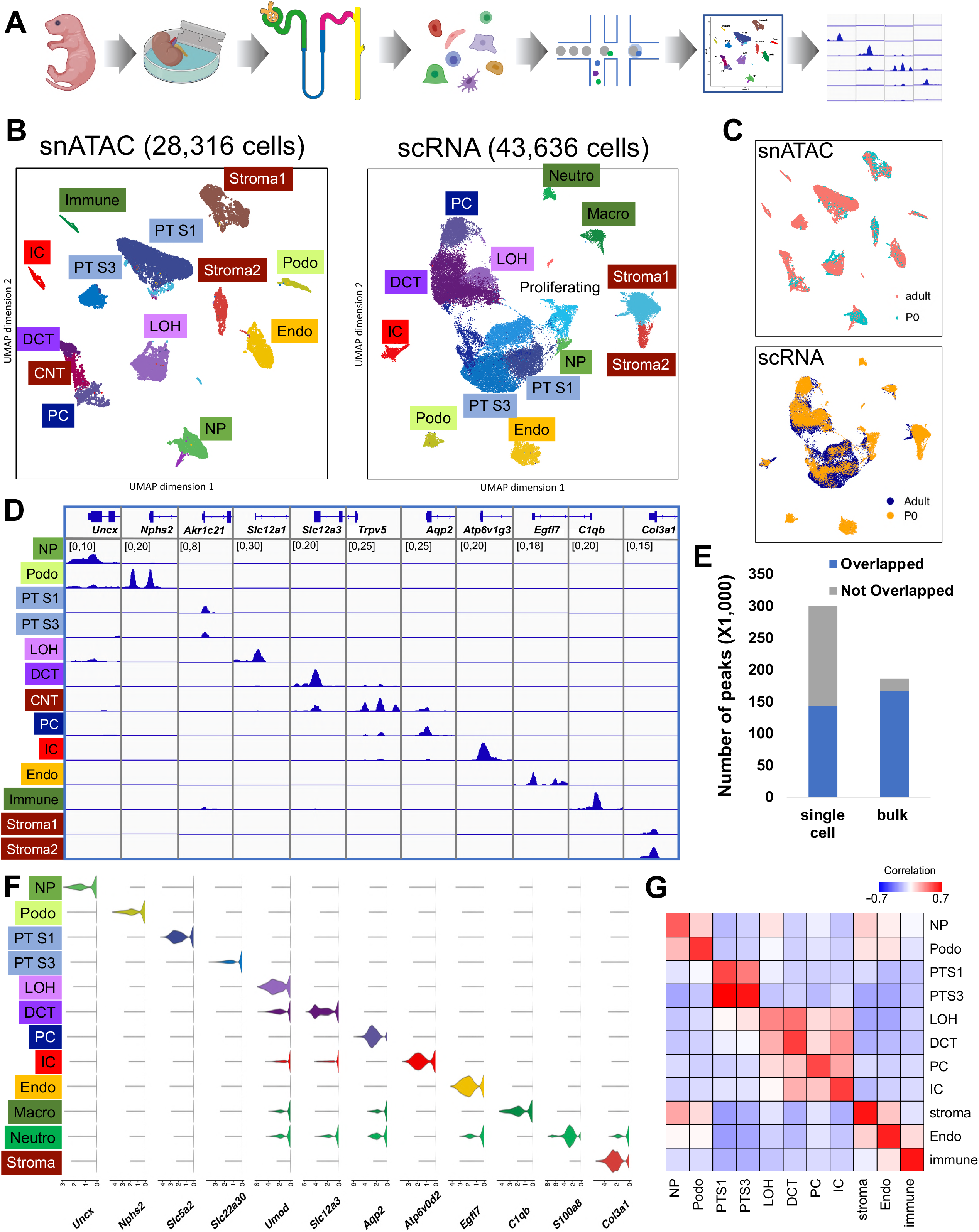
snATAC-seq and scRNA-seq identified major cell types in developing and adult mouse kidney. (A) Schematics of the study design. Kidneys from P0 and adult mice were processed for snATAC-seq and scRNA-seq followed by data processing and analysis including cell type identification and peak calling. (B) UMAP embeddings of snATAC-seq data and scRNA-seq data. Using marker genes, cells were annotated into nephron progenitors (NP), collecting duct intercalated cells (IC), collecting duct principal cells (PC), proximal tubule segment 1 and 3 (PT S1 and PT S3), loop of Henle (LOH), distal convoluted tubules (DCT), stromal cells (Stroma), podocytes (Podo), endothelial cells (Endo) and immune cells (Immune). In scRNA-seq data, the same cell types were identified, with an additional proliferative population and immune cells were clustered into neutrophils and macrophages. (C) UMAP embeddings of snATAC-seq and scRNA-seq data colored by P0 and adult batches. (D) Genome browser view of read density in each snATAC-seq cluster at cell type marker gene transcription start sites. We used Uncx for nephron progenitors, *Nphs2* for podocytes, *Akr1c21* for proximal tubules, *Slc12a1* for loop of Henle, *Slc12a3* for distal convoluted tubule, *Trpv5* for connecting tubule, *Aqp2* for collecting duct principal cells, *Atp6v1g3* for intercalated cells, *Egfl7* for endothelial cells, *C1qb* for immune cells and *Col3a1* for stroma. Additional marker gene examples are shown in **Figure S1h**. (E) Comparison of peaks identified from snATAC-seq data and bulk ATAC-seq data. Peaks that are identified in both datasets are colored blue, and peaks that are dataset-specific are grey. (F) Violin plots showing cell type-specific gene expression in scRNA-seq data. With the exception of proximal tubule, the same marker genes as in snATAC-seq data were used (*Slc5a2* and *Slc22a30* for proximal tubule S1 and S3, respectively). (G) Correlation between snATAC-seq gene activity scores and gene expression values in P0 data. The correlation of the adult dataset is shown in **Figure S1p**.

We next revealed cell type annotations from the open chromatin information. After conducting stringent filtering of the number of barcodes, promoter ratio and mitochondria ratio (**Methods, Figure S1d**), we kept 28,316 cells across the samples (**Figures 1b, S1f-g**). Cells were then clustered using snapATAC ^16^, which binned the whole genome into 5 kb regions and used diffusion map and principal component analysis for dimension reduction (**Methods**). Prior to clustering, we used Harmony ^17^, an iterative batch correction method, to correct for variability across samples. Using batch-corrected low dimensional embeddings, we clustered all cells together and retained 13 clusters, all of which had consistent representation across the number of peaks, samples and read depth profiles (**Figures 1b, S1e-g**). As expected, some clusters such as nephron progenitors and stromal cells were enriched in the developing kidney (P0).

In order to identify the cell type-specific open chromatin regions, we conducted peak calling using MACS2 ^18^ on each cell type separately. The peaks were then merged to obtain a comprehensive open chromatin set. We found that the single nuclei open chromatin set showed good concordance with bulk ATAC-seq samples, with most of the peaks in bulk ATAC-seq data captured by the single nuclei data. On the other hand, single nuclei chromatin accessibility data showed roughly 50% more accessible chromatin peaks (total of 300,693 peaks) than the bulk ATAC-seq data (**Figure 1e, Methods**), indicating that the snATAC-seq data was particularly powerful in identifying open chromatin areas that are accessible in single cell types.

To determine the cell types represented by each cluster, we examined chromatin accessibility around the TSS and gene body regions of the cognate known cell type-specific marker genes ^19^. Based on the accessibility of the known marker genes, we identified clusters representing nephron progenitors, endothelial cells, podocytes, proximal tubule segment 1 and segment 3 cells, loop of Henle, distal convoluted tubule, connecting tubule, collecting duct principal cells, collecting duct intercalated cells, stromal and immune cells (**Figure 1b**). **Figures 1d and S1h** show chromatin accessibility information for key cell type marker genes, such as *Uncx* and *Cited1* for nephron progenitors, *Nphs1* and *Nphs2* for podocytes, *Akr1c21* for both segments of proximal tubules, *Slc34a1* and *Slc5a2* for segment 1 of proximal tubules, *Kap* for segment 3 of proximal tubules, *Slc12a1* and *Umod* for loop of Henle, *Scl12a3* and *Pvalb* for distal convoluted tubule, *Trpv5* for connecting tubule, *Aqp2* and *Fxyd4* for principal cells, *Atp6v1g3* and *Atp6v0d2* for intercalated cells, *Egfl7* for endothelial cells, *C1qb* for immune cells and *Col3a1* for different types of stromal cells, respectively ^19^.

To understand cell type-specific gene expression changes, we also generated a single cell RNA sequencing (scRNA-seq) atlas for mouse kidney samples at the same developmental stages. The single cell transcriptome profiles of P0 and adult mouse kidneys were derived and processed as described in **Methods**. Rigorous quality control yielded a set of 43,636 single cells (**Figures 1b, S1i**). Quality control metrics such as gene counts, UMI counts and mitochondrial gene percentage along with batch correction results are shown in **Figures S1j-m**. By unbiased clustering ^20^ we obtained 17 distinct cell populations in the combined P0 and adult mouse datasets (**Figure S1i**). On the basis of marker gene expression, we identified kidney epithelial, immune and endothelial cells (**Figures 1f, S1n-o**), closely resembling the clustering obtained from snATAC-seq analysis. We then conducted differential expression analysis on the clusters and identified key marker genes for each cell type (**Supplemental Table 1**).

To compare the consistency between cluster assignment in the snATAC-seq data and the scRNA-seq data, we next derived a gene activity score for the top 3,000 highly variable genes in each snATAC-seq cluster and computed the Pearson’s correlation coefficient between each snATAC cluster and scRNA cluster (**Methods**). This analysis indicated good concordance between the two datasets (**Figures 1g, S1p**). While the correlation between gene expression and inferred gene activity score was high, we noted some differences in cell proportions, which was mostly related to the sample preparation-induced cell drop-out (**Figures S1g, i**). Consistent with previous observations that single cell preparations better capture immune cells than single nuclear preparations ^21^, we noted that the immune cell repertoire was limited in the snATAC-seq dataset; on the other hand, stromal cells were better captured by the nuclear preparation.

Finally, to allow the interactive use of this dataset by the community, we not only made the raw data available but also the processed dataset via our searchable website (susztaklab.com/developing_adult_kidney/snATAC/ for snATAC-seq data, susztaklab.com/developing_adult_kidney/scRNA/ for scRNA-seq data, and susztaklab.com/developing_adult_kidney/igv/ for IGV view of peak tracks). For example, here we show the chromatin accessibility landscape of *Ace2*, which is of major interest currently due to the COVID19 epidemic. We can observe an open chromatin region around the transcription start site of *Ace2* only in proximal tubules, which is consistent with its expression in proximal tubules (**Figure S1q**).

### Characterization of the cell type-specific regulatory landscape

To characterize different genomic elements captured by snATAC-seq data, we first stratified the genome into promoters, exons, 5’ and 3’ untranslated regions, introns, and distal regions using the GENCODE annotation ^22^ (**Methods**). We noticed that concordant with bulk ATAC-seq data, most peaks in snATAC-seq data were in regions characterized as distal elements or introns, (**Figure S2a**) and relatively small portions (<10%) were in promoter or 5’ untranslated regions. The genomic elements proportion was stable across developmental stages. In addition, almost half of the open chromatin peaks overlapped with P0 or adult H3K27Ac ChIP-seq signals (**Figure S2b**), indicating the contribution of enhancer regions to accessible chromatins.

To study the open chromatin heterogeneity in different cell types, we derived a cell type-specific accessible chromatin landscape by conducting pairwise Fisher’s exact test for each peak between every cluster (Benjamini-Hochberg adjusted *q* value ≤ 0.05, **Methods**). In total, we identified 60,684 differentially accessible open chromatin peaks (DAPs) across the 13 cell types (**Supplemental Table 2, Figure 2a**). Among these peaks, most showed high specificity for a single cluster. However, we noticed overlaps between the S1 and S3 proximal tubule segments-specific peaks, as well as between the loop of Henle and distal convoluted tubule segments, which is consistent with their biological similarities. In addition to the cell type-specific peaks, we also found some cell-type independent open chromatin areas (present across nephron progenitors, podocytes, proximal tubule and loop of Henle cells), likely consist of basal housekeeping genes and regulatory elements. (**Figures 2a, S2c**).

**Figure 2.**
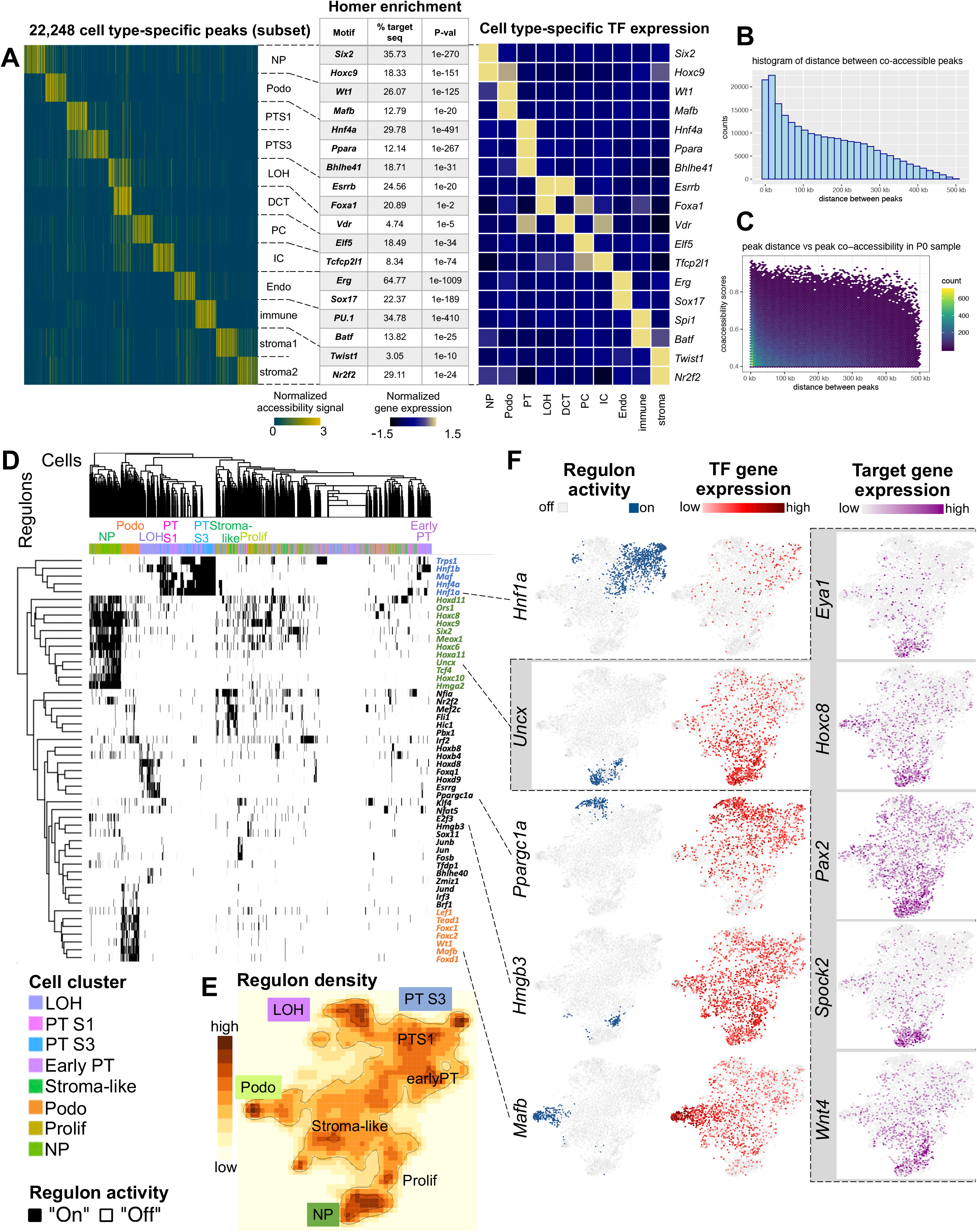
Cell type-specific gene regulatory landscape of the mouse kidney. (A) Left panel: Heatmap showing examples of the cell type-specific differentially accessible peaks (DAPs) (yellow: open chromatin, blue: closed chromatin) (full results are shown in **Supplemental Table 2**). Middle panel: Examples of cell type-specific motif enrichment analysis using Homer. (full results are shown in **Supplemental Table 3**). Right panel: TF expression z score heatmap that corresponds to the motif enrichment in each cell type. (B) Histogram showing the distribution of distances between Cicero-inferred correlated peaks in the P0 samples. The number of co-accessible peaks decreases with increasing distance. (C) Density plot showing the distribution of peak co-accessibility scores and distance between peaks. Although the number of peaks decreases with increasing distances, the co-accessibility score distribution remains relatively stable. (D) Regulon activity heatmap. Each column represents a single cell, colored by cluster assignment and ordered by hierarchical clustering; each row represents binarized regulon activities (“on-black”, “off-white”) and ordered by hierarchical clustering. (E) tSNE representation of regulon density as a surrogate for stability of regulon states, as inferred by SCENIC algorithm. (F) tSNE depiction of regulon activity (“on-blue”, “off-grey”) and TF gene expression (red scale) of exemplary regulons for proximal tubule (*Hnf1a*), nephron progenitors (*Uncx*), loop of Henle (*Ppargc1a*), proliferating cells (*Hmgb3*) and podocytes (*Mafb*). Examples of target gene expression of the *Uncx* regulon (*Eye1*, *Hoxc8*, *Pax2*, *Spock2* and *Wnt4*) are shown in purple scale. Expression of target genes of *Hnf1a*, *Ppargc1a*, *Hmgb3* and *Mafb* is shown in **Figure S3d**.

We noticed that many genes had strong cell type-specific DAPs at their TSS. Other genes, however, had accessible chromatin at their TSS in multiple cell types. For example, *Umod*, the loop of Henle-specific marker gene, showed accessible chromatin at its TSS at multiple tubule cell types (**Figure S2d, S1h**). Rather than with its TSS, cell type-specific chromatin accessibility of *Umod* strongly correlated with an upstream open chromatin peak, which is likely an enhancer region, indicated by the H3K27Ac ChIP-seq signal (**Figure S2d**). In addition, we noticed the enrichment of intronic regions and distal elements (**Figures S2e-f**) in cell type-specific DAPs, indicating their role in cell type-specific gene regulation.

These observations motivated us to study cis-regulatory elements using the snATAC-seq data and scRNA-seq data. We reasoned that a subset of the cell type-specific cis-regulatory elements should regulate cell type-specific gene expression in cis. Inspired by Zhu et al. ^23^, we aligned DAPs and differentially expressed genes from our snATAC-seq and scRNA-seq datasets, and inferred the putative regulatory peaks by their proximity (**Methods**). Such cis-regulatory elements predictions were confirmed by comparing with cis-regulatory elements inferred previously ^24^, as we recapitulated roughly 20% of elements from their analysis. In addition, our analysis was able to identify several known enhancers such as for *Six2* and *Slc6a18* ^24,25^ (**Figure S2g**).

To quantify the contribution of cis-regulatory elements, we analyzed peak co-accessibility patterns using Cicero ^26^. By using a heuristic co-accessible score 0.4 as a cutoff, we identified 232,380 and 206,701 cis-regulatory element links in the P0 and adult data, respectively. Some of these are likely promoter-enhancer regulatory units. Among these co-accessible elements, only 74,694 were common in P0 and adult kidneys, while most were developmental stage-dependent. While this observation needs further experimental validation, it highlights dynamic changes in gene regulation during development.

Another long-standing question has been to define the range of distances between interacting cis regulatory elements (such as enhancer-enhancer or enhancer-promoter). To this end, we explored the distances between co-accessible peaks using the Cicero output. We found that the number of co-accessible peaks decreased with increasing distance between open chromatin regions (**Figure 2b**). 43% of the co-accessible peaks were within 100 kb distance, however, the strength of association did not diminish with increasing distance. Even peaks that were as far as 500 kb apart showed high (0.8) co-accessibility scores (**Figure 2c**). Overall, the median distance between cis regulatory elements was relatively large (125.4 kb) and the number of interactions decreased with increasing distance, however, the strength of association did not change with increasing distance.

Given the complex interaction between genomic regions, we next looked into identifying key TFs that occupy the cell type-specific open chromatin regions. Until now, information on cell type-specific TFs in the kidney has been scarce. Therefore, we performed motif enrichment analysis on the cell type-specific open chromatin regions using HOMER ^27^. HOMER was designed as a differential motif discovery algorithm that scores motifs by computing enrichment of motif sequences in target compared to a reference set. To reduce false discovery, we focused on the known motifs. The full list of cell type-specific TF binding motifs is shown in **Supplemental Table 3**. Since several TFs have identical or similar binding sequences, we next correlated motif enrichment with scRNA-seq TF expression. Using this combined motif enrichment and gene expression approach, we have defined the mouse kidney cell type-specific TF landscape. Examples include *Six2* and *Hoxc9* in nephron progenitors, *Wt1* and *Mafb* in podocytes, *Hnf4a*, *Ppara*, and *Bhle41* in proximal tubules, *Esrrb* and *Foxa1* in loop of Henle, *Vdr* in distal convoluted tubule, *Elf5* in principal cells, *Tcfcp2l1* in intercalated cells, *Erg* and *Sox17* in endothelial cells, *Spi1* and *Batf* in immune cells, and *Twist1* and *Nr2f2* in stromal cells (**Figures 2a, S2h**).

In order to study the putative target genes of TFs, we examined TF regulon activity using Single-Cell rEgulatory Network Inference and Clustering (SCENIC) ^28^. SCENIC was designed to reveal TF-centered gene co-expression networks. By inferring a gene correlation network followed by motif-based filtration, SCENIC keeps only potential direct targets of each TF as modules (regulons). The activity of each regulon in each cell was quantified and then binarized to “on” or “off” based on activity distribution across cells (**Methods**). SCENIC was also able to conduct clustering based on the regulon states of each cell. SCENIC results (**Figures 2d-f**) indicated strong enrichment in *Trps1*, *Hnf1b*, *Maf*, *Hnf1a*, and *Hnf4a* regulon activity in proximal tubules, *Hmga2*, *Hoxc6*, *Hoxd11*, *Meox1*, *Six2*, *Tcf4*, and *Uncx* in nephron progenitors, *Esrrg*, and *Ppargc1a* in loop of Henle, *Hmgb3* in proliferating cells and *Foxc1*, *Foxc2*, *Foxd1*, *Lef1*, and *Mafb* in podocytes, respectively. While the expression of several of these TFs was relatively low and was further exacerbated by transcript drop-outs, many TFs did not show strong cell type enrichment. The regulon-based analysis, however, showed a very clear enrichment. SCENIC also successfully reported multiple downstream target genes. The full list of regulons and their respective target genes can be found in **Supplemental Table 4,** scaled and binarized regulon activity is also available in **Supplemental Table 5.** Examples of regulon activity, corresponding TF expression, and target gene expression are depicted in **Figures 2f, S2i**. For example, TFs such as *Eya1*, *Hoxc8*, *Hoxc9*, *Pax2*, *Spock2* and *Wnt4* are important downstream targets within the regulon of nephron progenitor-specific TF *Uncx*, indicating an important transcriptional hierarchy of nephron development ^29^. Finally, comparing the number of cell type-specific TFs reported by HOMER and SCENIC to the number of cell type-specific TFs among DEGs from RNA expression data, it became evident that cis-regulatory analysis in both snATAC-seq and scRNA-seq datasets yielded significant benefits in discovering the TF-regulatory network over analyzing transcript data alone (**Figures S2j-k**).

In summary, we generated a comprehensive atlas for the cell type-specific regulatory elements and TF-centered regulatory network.

### The regulatory trajectory of nephron progenitor differentiation

All cells in the body differentiate from the same genetic template. Cell type-specific chromatin opening and closing events associated with TF binding changes set up the cell type-specific regulatory landscape resulting in cell type specification and development. We found that closing of open chromatin regions was the predominant event during the nephron progenitor differentiation (**Figure S2a**). We then evaluated the cellular differentiation trajectory in the snATAC-seq and scRNA-seq datasets (**Methods**). We identified multiple nephron progenitor sub-groups (**Figures 3a-b**), which will need to be carefully mapped to prior gene expression- and anatomical location-driven nephron progenitor sub-classification. Consistently, across both data modalities, we identified that the podocyte precursors differentiated early from the nephron progenitor pool (**Figures 3a-b**). The tubule cell trajectory was more complex with a shared intermediate stage and later differentiation into proximal tubules and distal tubules/loop of Henle (**Figures 3a-b, S3a-c**). We also integrated snATAC-seq and scRNA-seq data to obtain a single trajectory (**Methods**). The cell types in this dataset were correctly mapped and the trajectory resembled the path observed in individual analyses of the scRNA and snATAC datasets (**Figures S3e-g**). The robustness of developmental trajectories was further supported by obtaining similar results when performing RNA velocity analysis using Velocyto ^30^ (**Figure S3d**) and by comparing with previous human and mouse kidney developmental studies ^9,13,14^.

**Figure 3.**
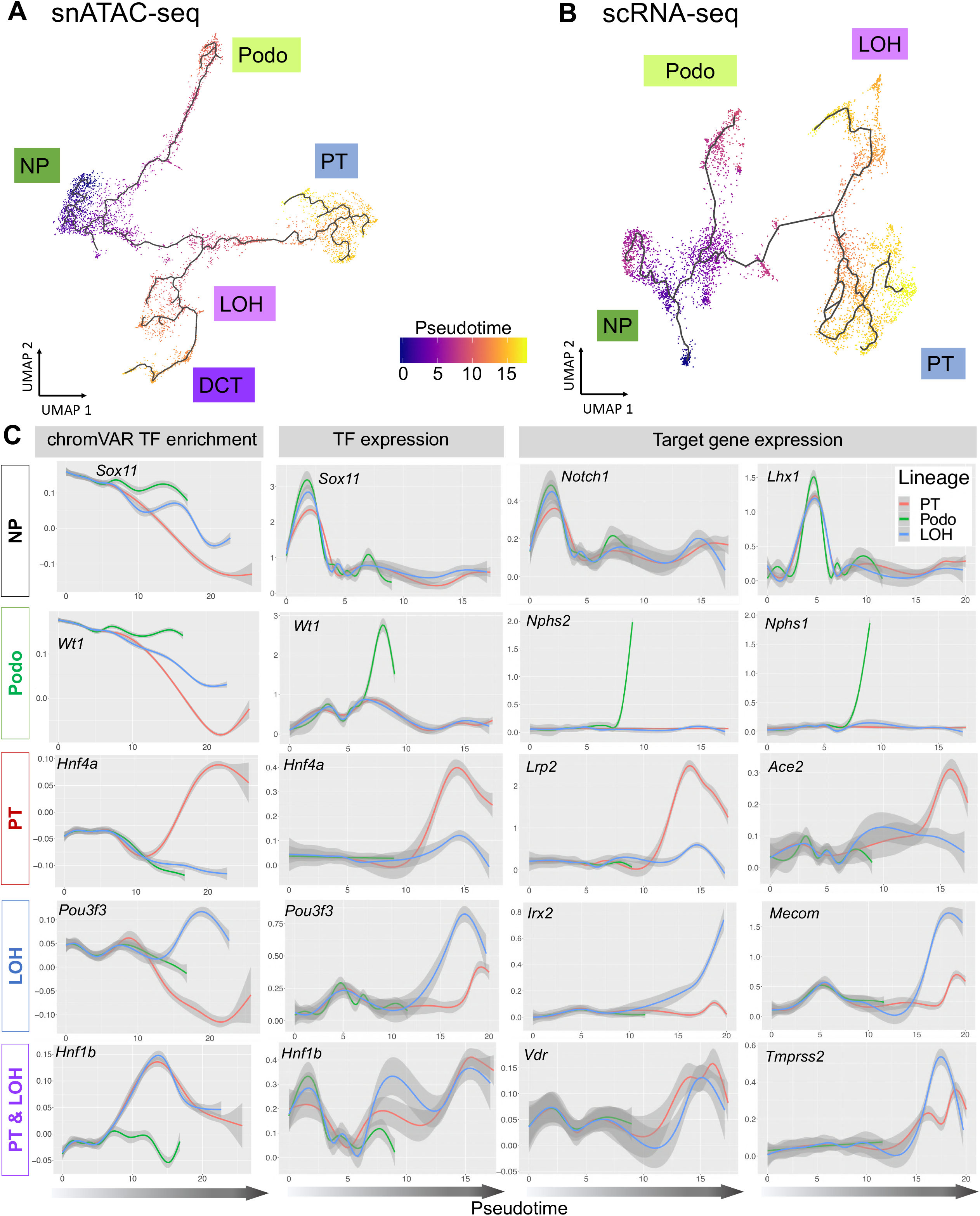
The cellular trajectory of nephron progenitor differentiation. (A) UMAP representation of snATAC-seq nephron progenitor differentiation trajectory towards podocytes, proximal tubule, loop of Henle and distal convoluted tubule, respectively, as inferred by Cicero. Cells are colored by pseudotime. (B) AP representation of scRNA-seq nephron progenitor differentiation trajectory towards podocytes, proximal tubule and loop of Henle, respectively, as inferred by Monocle3. Cells are colored by pseudotime. (C) Pseudotime-dependent chromatin accessibility and gene expression changes along the proximal tubule (red), podocyte (green) and loop of Henle (blue) cell lineages. The first column shows the dynamics of chromVAR TF enrichment score, the second column shows the dynamics of TF gene expression values and the third and fourth column represent the dynamics of SCENIC-reported target gene expression values of corresponding TFs, respectively. Additional examples are given in **Figure S3e**.

Building on both the SCENIC-generated gene regulatory network and the robust differentiation trajectories of the snATAC-seq and scRNA-seq datasets, we next aimed to understand chromatin dynamics, identify TFs and driver pathways for cell type specification and differentiation. To this end, we first determined variation in chromatin accessibility along the 3 differentiation trajectories using ChromVAR ^31^. ChromVAR estimates the accessibility dynamics of motifs in snATAC-seq data (**Methods**). We observed three different patterns when analyzing genes of interest (**Figure 3c**): *1) Decrease of TF motif accessibility in all lineages.* For example, *Sox11* motif enrichment score was high in nephron progenitor cells at the beginning of all 3 trajectories. It then decreased in all 3 lineages in parallel, underlining the role of *Sox11* in early kidney development. Several other TFs followed this pattern such as *Six2* and *Sox9. 2) Cell type-specific maintenance of chromatin accessibility with advancing differentiation.* We observed that chromatin accessibility for the *Wt1* motif was high initially but declined in proximal tubule and loop of Henle lineages, while its expression increased in the podocyte lineage. This is consistent with the important role of *Wt1* in nephron progenitors and podocytes ^32,33^. Other TFs that followed this pattern include *Foxc2* and *Foxl1*. *3) A de novo increase in chromatin accessibility with cell type commitment and advancing differentiation.* For example, the chromatin accessibility of *Hnf4a* and *Pou3f3* motif increased in proximal tubule and loop of Henle trajectories, respectively, coinciding with the cellular differentiation program ^34^. A large number of TFs followed this pattern such as *Mafb* (in podocytes), *Hnf4a* and *Hnf1a* (in proximal tubule), *Hnf1b* (in both proximal tubule and loop of Henle) as well as *Esrrb* and *Tfap2b* (in loop of Henle).

Next, we correlated changes in chromatin accessibility-based TF motif enrichment with TF expression and their respective target genes along Monocle-generated trajectories. To this end, we used the scRNA-seq differentiation trajectories to find TFs and target genes differentially expressed over pseudotime (**Supplemental Table 6)**. We also noticed a good concordance of time-dependent changes of TF and target gene expression along with TF motif enrichment, including the lineages for podocytes (e.g., *Foxc2*, *Foxl1*, *Mafb*, *Magi2*, *Nphs1*, *Nphs2*, *Plat*, *Synpo*, *Thsd7a*, *Wt1*, and *Zbtb7c*), proximal tubule (e.g., *Ace2*, *Atp1a1*, *Dab2*, *Hnf1a*, Hnf4a, *Hsd17b2*, *Lrp2*, *Maf*, *Slc12a3*, *Slc22a12*, *Slc34a1*, and *Wnt9b*), loop of Henle (e.g.,*Cyfip2*, *Cytip*, *Esrrb*, *Esrrg*, *Irx1*, *Irx2*, *Mecom*, *Pla2g4a*, *Pou3f3*, *Ppargc1a*, *Stat3*, *Sytl2*, *Tfap2b*, *Thsd4*, and *Umod*), as well as for both proximal tubule and loop of Henle (e.g., *Bhlhe40*, *Hnf1b*, and *Tmprss2*), respectively (**Figures 3c, S3i**). Most interestingly, we noticed two distinct patterns of how gene expression was related to chromatin accessibility. While gene expression of TFs increased over pseudotime, its corresponding motif accessibility either increased in parallel (such as *Hnf4a* and *Pou3f3*) or maintained in a lineage-specific manner (such as *Wt1*). This might indicate different regulatory mechanisms during differentiation.

We next aimed to interrogate the stage-dependent chromatin dynamics along the identified differentiation trajectory. The differentiation trajectory was binned into 15 developmental steps based on the lineage specification (**Figures S3b-c**). These stages were labeled as NP (nephron progenitor), IM (intermediate cells), Podo (podocytes), PT (proximal tubule), LOH (loop of Henle), and DCT (distal convoluted tubule), however, this designation will need to be matched with prior cell marker-based annotations. To study the chromatin opening and closing, we conducted differential chromatin accessibility analysis between subsequent stages. To understand the biological processes controlled by the epigenetic changes, we examined the nearest genes and performed functional annotation (**Methods**). We found that open chromatin profiles were relatively stable in the early precursor stages such as NP1 to NP3, with fewer than 70 DAPs identified (**Supplemental Table 7, Figure S4a**). The podocyte differentiation branch was associated with marked increase in the number of DAPs, (796 DAPs between NP3 and Podo1). This mainly represented the closing of chromatin areas around nephron progenitor-specific genes such as *Osr1*, *Gdnf*, *Sall1*, *Pax2* and opening of areas around podocyte-specific genes and key TFs such as *Foxc2* and *Efnb2*, both of which are validated to be important for early podocyte differentiation ^35,36^. At later stages, there was a strong increase in expression of actin filament-based processes and a significant decrease in *Notch* and *Ctnnb1* in the podocyte lineages (**Supplemental Table 8**). Fewer chromatin closing events were observed (234 DAPs) between NP3 and intermediate cells 1 (IM1), mainly associated with closing of the chromatin around *Osr1* and opening around tubule cell-specific TFs such as *Lhx1* and *Pax3* (**Figure 4**). The decrease in *Six2* expression only occurred at the IM2 stage, at which we also observed an increase in tubule specification genes such as *Hnf1a*. Gene ontology results from the 820 up-regulated peaks between PT1 and IM2 showed enrichment associated with typical proximal tubule functions including sodium-dependent phosphate transport, maintenance of osmotic response in the loop of Henle and active sodium transport in the distal convoluted tubule (**Figure S4a**, the full list can be found in **Supplemental Tables 7, 8 and 9)**.

**Figure 4.**
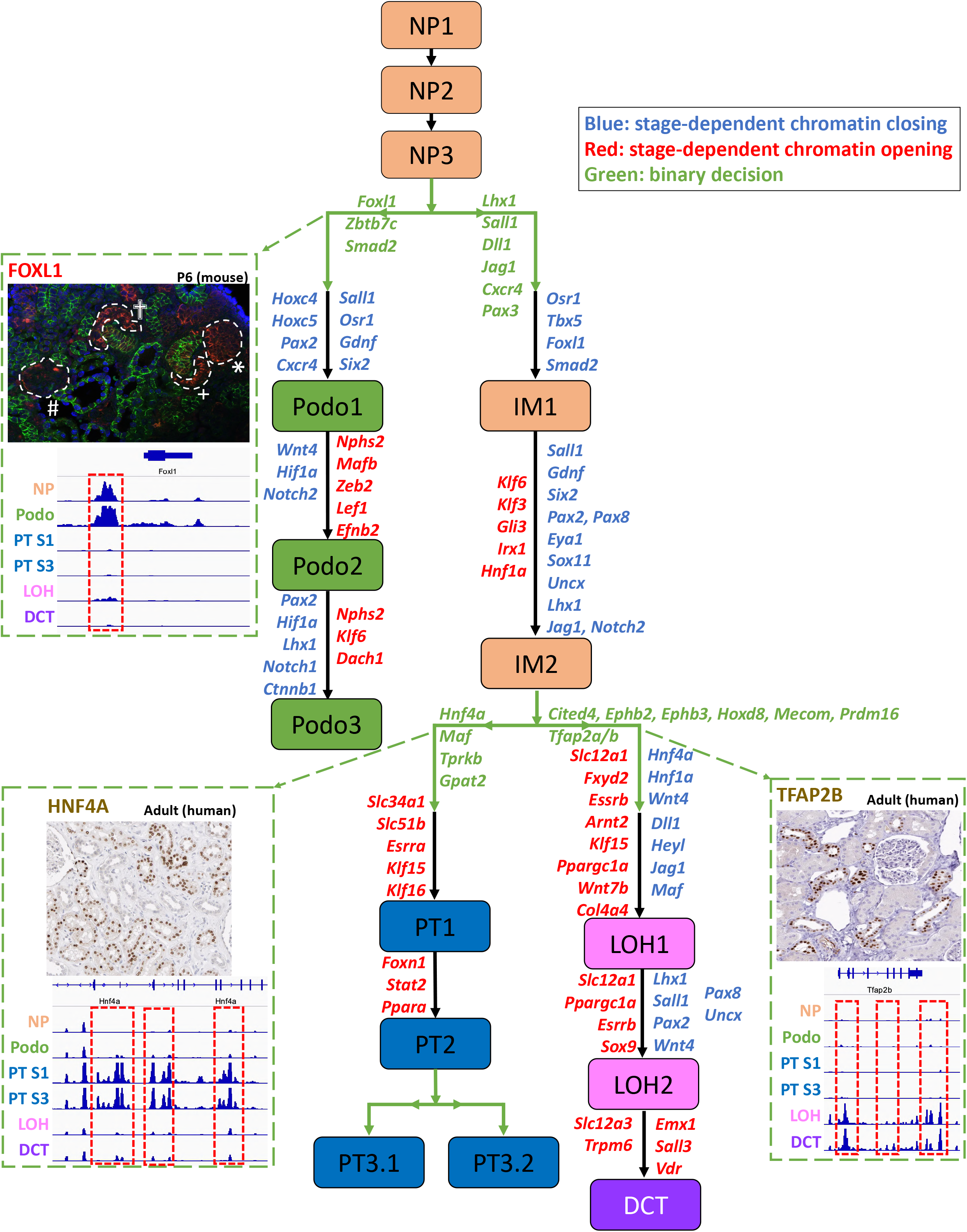
Chromatin dynamics of nephron progenitor differentiation. Di-graph representing cell type and lineage divergence, as derived from Cicero trajectory inference. Nephron progenitors (NP), podocytes (Podo), intermediate stage (IM), proximal tubule (PT), loop of Henle (LOH) and distal convoluted tubule (DCT) are connected with their developmental precursor stages and represented by ascending numbering. Arrows represent cell differentiation along respective trajectories. Genes listed next to the trajectories were derived from analyzing gene enrichment of differentially assessible peaks (DAPs) between two stages. Genes colored red were derived from the opening DAPs between two stages, genes colored blue were derived from the closing DAPs between two stages, and genes colored green were derived from opening DAPs between two branches. Three important genes, *Foxl1*, *Hnf4a* and *Tfap2b* are shown along with their cell type-specific accessibility peaks and immunostaining results. Peaks that were open during the development of specific cell types are shown in red boxes. Immunofluorescence staining of fetal mouse kidney shows FOXL1 in red along cellular differentiation (from right to left) from early progenitor stage (asterisk) over comma-shaped (+) and S shaped bodies (cross) towards podocytes within primitive glomeruli (#). HNF4A and TFAP2B in human adult kidney samples (taken from the Human Protein Atlas, http://www.proteinatlas.org ^62^) are visualized by immunohistochemistry in brown.

In addition to analyzing changes along the trajectory, we also specifically examined cell-fate decision events. We studied the chromatin opening and closing during the first cell commitment event. We found that podocyte specification from nephron progenitors was associated with differential opening of *Foxl1*, *Zbt7c,* and *Smad2* in the podocyte lineage and *Lhx1*, *Sall1*, *Dll1*, *Jag1*, *Cxcr3* and *Pax3* in the other lineage, respectively. While the role of several TFs has been established for podocyte specification, the expression of *Foxl1* has not been described in the kidney until now (**Figure 4**). Our analysis pinpointed that four peaks in the vicinity of *Foxl1* were accessible only in podocyte lineage, which locate in +53,381 bp, +152,832 bp, +237,019 bp, and +268,550 bp of the *Foxl1* TSS, respectively. To confirm the expression of *Foxl1* in nephron progenitors and podocytes, we performed immunofluorescence studies on developing kidneys (E13.5, P0 and P6). Consistent with the computational analysis, we found strong expression of FOXL1 in nephron progenitors (E13.5). At later stages, it was present in comma and S shape body and finally in the glomerular podocytes (**Figure S4b**). While there was no expression within cells destined to become proximal tubule or loop of Henle cells, gene expression of *Foxl1* increased in cells along the podocyte trajectory (**Figure S4c**). While further experimental validation will be important, our study has illustrated the critical role of open chromatin state information and dynamics in cellular differentiation.

The intermediate cells (IM) gave rise to proximal and distal branches, representing the proximal tubules and the loop of Henle as well as distal convoluted tubule segments. The proximal tubule region was characterized by chromatin opening around *Hnf4a, Maf*, *Tprkb*, and *Gpat2*. The loop of Henle and distal convoluted tubule segments were remarkable for multiple DAPs in the vicinity of *Tfap2a*, *Tfap2b*, *Cited4*, *Ephb2*, *Ephb3*, *Hoxd8*, *Mecom*, and *Prmd16*, indicating a critical novel role for these TFs in distal tubule differentiation (**Figures 3c, 4, S4c**). Consistently, we saw a reduction in chromatin accessibility of *Six2* promoter and enhancers along all three trajectories (podocyte, proximal tubule and loop of Henle) (**Figure S4d**). There was also a decrease in expression of *Jag1* and *Heyl* in the distal loop of Henle segment, concordant with the putative role of Notch driving the proximal tubule fate ^37^ **(Supplemental Table 11)**. Another striking observation was that tubule segmentation and specification occurred early by an increase in chromatin accessibility around *Lhx1*, *Hnf1a* and *Hnf4a* and *Maf* for proximal tubule and *Tfap2b* for loop of Henle. Terminal differentiation of proximal tubule and loop of Henle cells was strongly linked to nuclear receptors that regulate metabolism, such as *Esrra* and *Ppara* in proximal tubules and *Esrra* and *Ppargc1a* in the loop of Henle segment, once more indicating the critical role of metabolism of driving gene expression and differentiation ^38^.

In summary, we reconstructed the developmental and differentiation trajectories of podocytes, proximal tubule and loop of Henle cells. We defined chromatin and gene expression dynamics and identified numerous putative TFs for kidney cell specification and differentiation.

### Stromal-to-epithelial communication is critical in the developing and adult kidneys

Previous studies indicated that the survival, renewal, and differentiation of nephron progenitors is largely regulated through its cross-talk with the adjacent ureteric bud ^39^. To investigate the complex cellular communication network, we used CellPhoneDB ^40^ to systematically infer potential cell-cell communication in the developing and adult kidney. CellPhoneDB provides a comprehensive database and a statistical method for the identification of ligand-receptor interactions in scRNA-seq data. Analysis of our scRNA-seq dataset indicated that the number of cell-cell interaction pairs was larger in developing kidney compared to the adult kidney (**Figure 5a**). In the developing kidney, the stroma showed the greatest number of interactions among all cell types, coinciding the well-known role of epithelial-stromal interactions in driving kidney development. Of the identified interactions, many were related to stroma-secreted molecules such as collagen 1, 3, 4, 6, and 14 (**Figure 5b**). Furthermore, the stroma seemed to interact with most cell types, such as podocytes and different tubule cells. Interestingly, the nephron progenitor cluster showed important ligand-receptor interaction between *Fgf1*, *Fgf8* as well as *Fgf9* and the corresponding receptor *Fgfr1*, which is consistent with the well-known role of FGF signaling in kidney development ^41^. Of the manifold identified interactions in the fetal kidney, stromal interaction and the VEGF-involving interaction remained significant in the adult data set, underscoring the importance of endothelial-to-epithelial communication.

**Figure 5.**
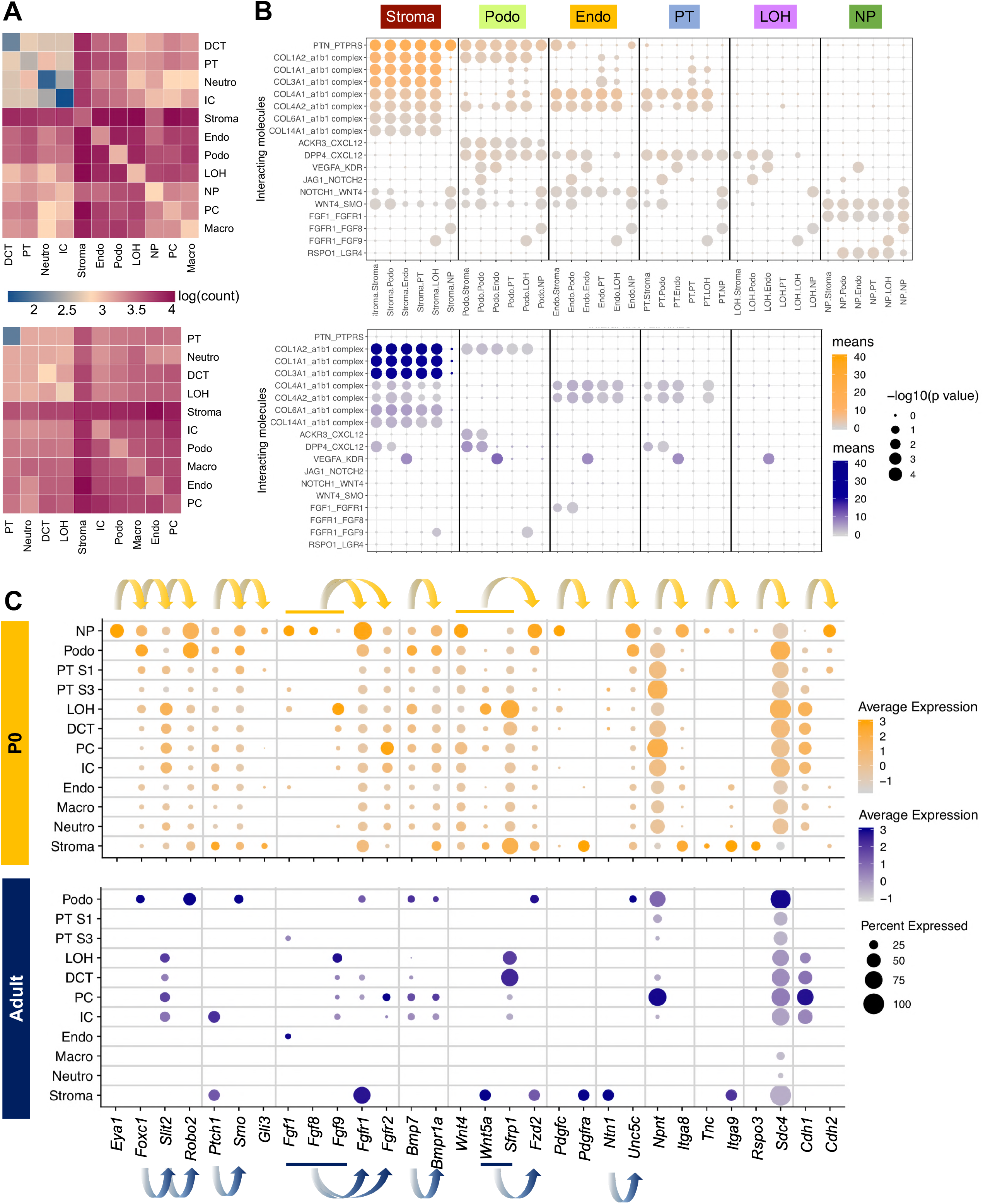
Cell-cell communication analysis in the developing and adult mice highlighted the critical role of stroma in driving cell differentiation. (A) Heatmaps showing the number of cell-cell interactions in the scRNA-seq dataset of P0 (top) and adult (bottom) kidneys, as inferred by CellPhoneDB. Dark blue and dark red colors denote low and high numbers of cell-cell interactions, respectively. (B) CellPhoneDB-derived measures of cell-cell interaction scores and p values. Each row shows a ligand-receptor pair, and each column shows the 2 interacting cell types, which is binned by cell type. Columns are sub-ordered by first interacting cell type into stroma, podocytes, endothelial cells, proximal tubule, loop of Henle and nephron progenitors. Color scale denotes the mean values for all the interacting partners, where mean value refers to the total mean of the individual partner average expression values in the interacting cell type pairs. Orange scale denotes P0, blue scale denotes adult. Dot size denotes corresponding p values of the permutation test. (C) Dot plots of RNA expression of important cell-cell communication candidates within the Gdnf-Ret, Sonic hedgehog, Fgf, Bmp, Wnt and other pathways in both P0 (top) and adult (bottom) kidney. Dot size denotes percentage of cells expressing the marker. Color scale represents average gene expression values, orange denotes P0, blue denotes adult. Arrows indicate ligand-receptor pairs.

We next individually examined the expression of several key pathways known to play important roles in kidney development, such as Gdnf-Ret, sonic hedgehog, FGF, Bmp, Wnt and others ^9^. Expression of these key ligand-receptor pairs showed strong cell type specificity (**Figure 5c**). For example, *Robo2* of the Gdnf-Ret pathway was expressed in nephron progenitors and in podocytes of P0 and adult kidney. Gdnf signaling through the Ret receptor is required for normal growth of the ureteric bud during kidney development ^42^ and the Slit2/Robo2 pathway is implicated with congenital kidney anomalies ^43^ and important for maintenance of podocyte foot process integrity ^44^. *Eya1*, however, is genetically upstream of *Gdnf* and acts as a positive regulator for its activation ^45^. Consistently, we noted distinct cell type specificity of *Eya1* expression only in nephron progenitors, which was also true for other important signaling molecules such as *Ptch1, Smo* and *Gli3* of the sonic hedgehog pathway. *Fgfr1* showed the highest expression in nephron progenitors as well as in fetal and adult stroma, underscoring the importance of FGF signaling for cell-cell interactions in both the developing and developed kidney. Most interestingly, some cell-cell interactions between specific cell types that were observed in fetal kidney were abrogated in adult kidney because of the loss of expression of either ligand or receptor, such as *Pdgfc* in nephron progenitor signaling to its receptor *Pdgfra* in stroma, *Npnt* from several epithelial cells signaling to *Itga8* in nephron progenitors, *Tnc-Itga9* signaling from nephron progenitors to stroma and *Rspo3* in stroma signaling to *Sdc4* in several epithelial cells. Because not much is known about some of these markers, the significance of these putative interactions requires further investigation. For example, *Rspo3* has been implicated in nephron progenitor-associated interactions during nephrogenesis ^10^. Mutations in the *Itga8* gene are known to cause isolated congenital anomalies of kidney and urinary tract in humans ^43^ and *Pdgfra* has been regarded as a commitment marker in kidney differentiation ^10^.

In summary, we inferred cell-cell interactions in the developing and adult kidneys and found the critical role of stromal-epithelial interactions in the developing kidney.

### Single cell chromatin accessibility identified human kidney GWAS target regulatory regions, genes and cell types

Finally, we examined whether single cell level chromatin accessibility data can help identify cell and gene targets for human kidney disease development. GWAS have been exceedingly successful in identifying nucleotide variations associated with specific diseases or traits. However, more than 90% of the identified genetic variants are in the non-coding region of the genome. Initial epigenome annotation studies indicated that GWAS hits are enriched in tissue-specific enhancer regions. As there are many different cell types in the kidney with differing function, understanding the true cell type specificity of these enhancers is critically important. Here, we reasoned that single cell accessible chromatin information could be extremely useful to identify the cell type-specific enhancer regions and thereby the target cell type for the GWAS hits, however, such maps have not been generated for the human kidney. We combined three recent kidney disease GWAS ^46–48^, and obtained 26,637 single nucleotide polymorphisms (SNPs) that passed genome-wide significance level of which we retained 7,923 after lift-over from human to mouse.

By overlapping the kidney disease-associated SNPs with peaks called by snATAC-seq data, we found highly specific accessibility among different cell types (**Figure S5**). We found that most of the peaks overlapped with nephron lineages, especially proximal tubules. The full table including nearest genes is provided in the **Supplemental Table 12**. However, as the causal GWAS variant is unknown, we conducted further investigation of kidney disease target genes and cell types using multi-omics mouse kidney data.

Specifically, we examined loci where functional validation studies reported conflicting results on target cell types and target genes (**Figure 6)**. The *SHROOM3* locus has shown a reproducible association with kidney function in multiple GWAS ^48^. However, previous functional follow-up studies have reported confusing and somewhat contradictory results. While one study indicated that the genetic variants were associated with an increase in SHROOM3 levels in tubule cells inducing kidney fibrosis ^49^, the other suggested that the variant was associated with lower SHROOM3 levels in podocytes resulting in chronic kidney disease development ^50^. We found an open chromatin (likely promoter) area in multiple cell types such as nephron progenitors, podocytes, loop of Henle, distal convoluted tubule, principal cells and intercalated cells (**Figure 6a**). We also identified intronic open chromatin areas only in nephron progenitors and podocytes that overlapped with the GWAS significant variants (**Figure 6a**). Consistent with the cis-regulatory open chromatin, the strongest expression of *Shroom3* was observed in podocytes and nephron progenitor cells. Expression of *Shroom3* in the adult bulk kidney was below our detection limit. To further understand the regulatory dynamics of this locus in the developing mouse kidneys, we examined gene expression and epigenome annotation data generated from bulk mouse kidney samples at different stages of development for H3K27ac and H3K4me1 in adult and fetal samples (**Figure 6b**). Interesting to note that the GWAS-significant SNP that showed strong nephron progenitor-specific enrichment also coincided with the *Six2* binding area. Finally, the Cicero-based co-accessible analysis connected the GWAS top variants, which located in an intronic enhancer region of *Shroom3*, with *Shroom3* exons, indicating that *Shroom3* is the likely target gene of the variant (**Figure 6a**).

**Figure 6.**
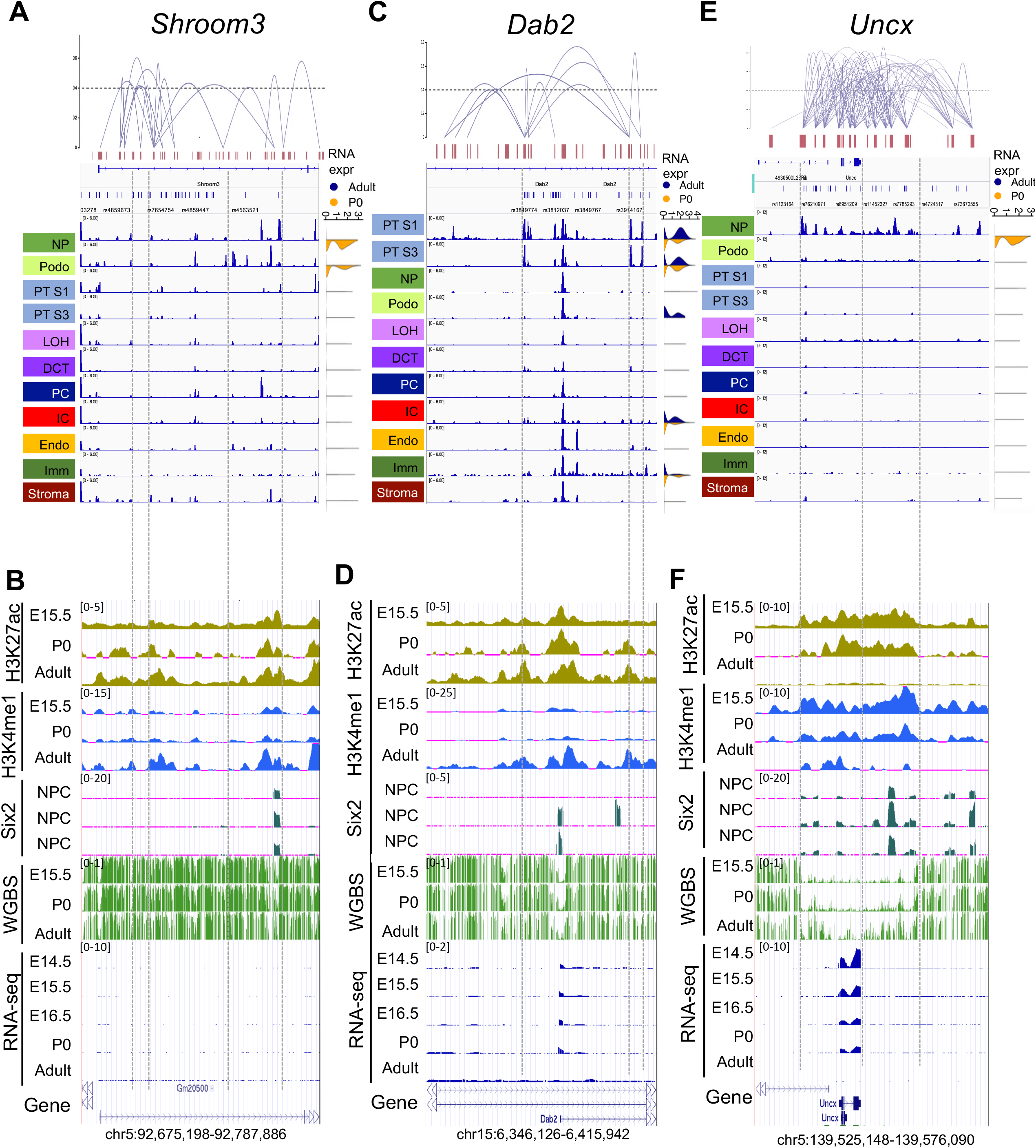
Single cell level chromatin accessibility highlighted human kidney GWAS target genes and cell types. (A, C, E) From top to bottom: Cicero-inferred co-accessibility of open chromatin regions in mouse orthologues of human *Shroom3*, *Dab2* and *Uncx* loci; Gene browser view of the single nucleotide polymorphisms within the regions; gene browser view of chromatin accessibility for nephron progenitors (NP), collecting duct intercalated cells (IC), collecting duct principal cell types (PC), proximal tubules segment 1 and 3 (PT S1 and PT S3), loop of Henle (LOH), distal convoluted tubule (DCT), stromal cells (stroma), podocytes (Podo), endothelial cells (Endo) and immune cells (Immune). Right subpanel shows violin plots of scRNA-seq gene expression in P0 (orange) and adult (blue) kidneys. (B, D, F) Whole kidney H3K27ac, H3K4me1 and Six2 ChIP-seq, whole genome bisulfate sequencing (WGBS) and RNA-seq data in E15.5, P0 and adult kidney samples.

Next, we analyzed the chromosome 15 GWAS region, where we identified some open chromatin regions that were uniformly open in all examined cell types. *Dab2* expression, on the other hand, strongly correlated with open distal enhancer regions in proximal tubule cells (**Figure 6c**). This is consistent with earlier publications indicating the role of proximal tubule-specific DAB2 playing a role in kidney disease development ^51^. Interestingly, while single cell analysis indicated an additional distal enhancer in intercalated cells, the GWAS-significant region coincided with the proximal tubule-specific enhancer region and showed strong coregulation (**Figure 6d**). Regulatory annotation of the developing kidney indicated strong enhancer marks in the adult but not in the fetal kidney.

Lastly, we examined the region around *Uncx*, for which reproducible association with kidney function was shown in multiple GWAS ^46,47^. Interestingly, the GWAS locus demonstrated a strong open chromatin region in nephron progenitors but not in any other differentiated cell types (**Figure 6e**). Consistently, in bulk chromatin accessibility data we only observed regulatory activity such as H3K27ac, H3Kme1 and we show *Six2*-binding at this locus in fetal kidneys. The locus did not show H3K27ac enrichment in the adult kidney, while H3K4me1 remained positive (**Figure 6f**). *Uncx* expression was strong in the fetal kidney samples, but we could not detect its expression in the adult kidney (**Figure 6e**). A closer view of these loci is shown in **Figure S6**.

These results indicate that variants associated with kidney disease development are located in regions with cell type- and developmental stage-specific regulatory activity and illustrate the critical role of snATAC-seq in defining target genes and target cell types for GWAS variants.

## Discussion

In summary, here we present the first cellular resolution open chromatin map for the developing and adult mouse kidney. Using this dataset, we identified key cell type-specific regulatory networks for kidney cells, defined the cellular differentiation trajectory, characterized regulatory dynamics and identified key driving TFs for nephron development, especially for the terminal differentiation of epithelial cells. Furthermore, our results shed light on the cell types and target genes for genetic variants associated with kidney disease development.

By performing massively parallel single cell profiling of chromatin state, we were able to define the key regulatory logic for each kidney cell type by investigating cis-regulatory elements and TF-target gene interaction. We found that most cell type-specific open chromatin regions are within distal regulatory elements and intronic regions. Our studies identified a massive amount of highly dynamic co-regulated peaks indicating the important correlation between distal regulatory elements and gene expression. Future studies will examine the relative contribution of promoters and enhancer openness in gene expression regulation. However, these studies highlight that both chromatin opening and looping are critical for gene regulation.

We also observed that the single cell open chromatin atlas was able to define more distinct cell types even in the developing kidney compared to scRNA-seq analysis. Given the continuous nature of RNA expression, it has been exceedingly difficult to dissect specific cell types in the developing kidney ^9,10,13^. In addition, it has been difficult to resolve the cell type origin of lowly expressed transcripts in scRNA-seq data. However, this is not the case for snATAC-seq data, which were able to capture the chromatin state irrespective of gene expression magnitude. There were several examples where accessible peaks were identified in specific cell types even for lowly expressed genes such as Shroom3.

We identified critical cell type-specific TFs by integrating multiple computational analyses. TF identification is challenging in scRNA-seq data since the expression of several cell type-specific TFs is low and some of them do not show a high degree of cell type-specificity ^52^. By extracting motif information, snATAC-seq data provides additional information for TF identification.

Together with regulon analysis, as implemented in SCENIC, we have identified several TFs as well as their target genes that are important for kidney development. Leveraging this newly identified cell type-specific regulatory network will be essential for future studies of cellular reprogramming of precursors into specific kidney cell types and for better understanding homeostatic and maladaptive regeneration.

Our studies revealed dynamic chromatin accessibility that tracks with renal cell differentiation. These states may reveal mechanisms governing the establishment of cell fate during development, in particular those underlying the emergence of specific cell types. We found a consistent and coherent pattern between gene expression and open chromatin information, where the nephron progenitors differentiated into two branches representing podocytes and tubule cells ^53^. We found that podocytes commitment occurred earlier, while tubule differentiation and segmentation appeared to be more complex. This podocyte specification correlated with the maintenance of expression of *Foxc2* and *Foxl1* expression in podocytes. While *Foxc2* has been known to play a role in nephron progenitors and podocytes, this is the first description of *Foxl1* in kidney and podocyte development. Our studies are consistent with recent observations from organoid models that recapitulated podocyte differentiation better than tubule cell differentiation ^54^. Our study also sheds light on tubule differentiation and segmentation. We confirmed the key role of *Hnf4a* in proximal tubules. We have identified a large number of new transcriptional regulators such as *Tfap2a* that seem to be critical for the distal portion of the nephron. Our data indicate that distal tubule differentiation is linked to the loop of Henle, a critically important observation needing further confirmation. Furthermore, the terminal differentiation of proximal tubule cells correlated with the increase in *Ppara* and *Esrra* expression, both of which are known regulators of oxidative phosphorylation and fatty acid oxidation ^38^. Loop of Henle differentiation strongly correlated with *Essrb* and *Ppargc1a* expression. These studies potentially indicate that cell specification events occur early and metabolism controls terminal differentiation of tubule cells ^55^. Impaired metabolic fitness of proximal tubules has been a key contributor to kidney dysfunction, explaining the critical association with tubule metabolism and function.

Furthermore, we show that single cell and stage level epigenome annotation is critical for the annotation of human GWAS. Most identified GWAS signals are in the non-coding region of the genome. Due to the linkage disequilibrium structure of the human genome, each GWAS locus contains a large number of variants, each passing genome-wide significance level ^56^. Furthermore, as these signals are often non-coding, the target gene and the target cell type remain unknown. While molecular quantitative trait locus studies and bulk epigenome annotation experiments have been important to define the molecular pathways leading to disease development from the identified signals, these methods have limited resolution, as cell type-specific enhancer regions cannot be identified by bulk analysis ^57^. Additionally, bulk molecular quantitative trait locus studies suffer from the same linkage disequilibrium problems as GWAS analyses ^58^. Our results indicate that multiple GWAS regions are conserved between mice and humans. Single cell open chromatin information enables not only the identification of affected cell types, but also the understanding of co-regulation of the open chromatin area. It is also able to highlight critical target genes. Performing single cell open chromatin analysis on human kidney tissue samples will be essential to further understand molecular pathways altered by genetic variants. Here we showed three important examples. We confirmed the role of *Dab2* and its specific expression in the proximal tubule during kidney disease development, as its implication therein has been shown in previous expression quantitative trait locus and bulk epigenome analysis experiments ^51^. Furthermore, we showed that the GWAS variants map only to those regions where chromatin is open exclusively in nephron progenitors, whereas chromatin becomes inaccessible as differentiation progresses during later stages, such as *Shroom3* and *Uncx*. This is an interesting and important novel mechanism, indicating that the altered expression of this gene might play a role in the development rewiring of the kidney. This mechanism is similar to genes associated with autism that are known to be expressed in the fetal but not in the adult stages ^59^ and highlights the critical role of understanding chromatin accessibility at multiple stages of differentiation.

While we have generated a large amount of high-quality data, this information will need further experimental validation, which is beyond the scope of the current manuscript. In addition, one needs to be aware of the limitations when interpreting different computational analyses, for example, the motif enrichment analyses such as implemented by HOMER, SCENIC, and chromVAR, are not able to distinguish between TFs with similar binding sites. Future high-throughput studies that analyze open chromatin and gene expression information from the same cells will be exceedingly helpful to correlate open chromatin and gene expression information along the differentiation trajectory ^24,60,61^.

In summary, our dataset provides critical novel insight into the cell type-specific gene regulatory network, cell differentiation program, and disease development.

## Acknowledgements

Work in the Susztak lab is supported by the NIH DK076077, DK087635, and DK105821. MSB is supported by a German Research Foundation grant (BA 6205/2-1).

## Author Contributions

KS and ZM designed and conceived the experiment. ZYM, JW, RS, and TA conducted the experiment. ZM conducted snATAC-seq bioinformatics analysis with advice from KS, HL, ML, and JK. MSB and ZM conducted scRNA-seq bioinformatics analysis with advice from KS. AMK and AYK conducted immunofluorescence staining with supervision from KHK. KS, ZM, and MSB wrote the manuscript and all authors edited and approved of the final manuscript.

## Methods

### Single cell RNA sequencing of P0 mice

1-day-old mouse neonate was decapitated with surgical scissors, 2 kidneys were harvested and minced into 1 mm^3^ pieces and incubated with digestion solution containing Enzyme D, Enzyme R and Enzyme A from Multi Tissue Dissociation Kit (Miltenyi, 130-110-201) at 37 °C for 15 min with agitation. Reaction was deactivated by adding 10% FBS, then solution was passed through a 40 μm cell strainer. After centrifugation at 1,000 RPM for 5 min, cell pellet was incubated with 500 μL of RBC lysis buffer on ice for 3 min. We centrifuged the cells at 1,000 RPM for 5 min at 4 °C and resuspended the cells in the buffer for further steps. Cell number and viability were analyzed using Countess AutoCounter (Invitrogen, C10227). The cell concentration was 2.2 million cells/mL with 92% viability. 10,000 cells were loaded into the Chromium Controller (10X Genomics, PN-120223) on a Chromium Single Cell B Chip (10X Genomics, PN-120262) and processed to generate single cell gel beads in the emulsion (GEM) according to the manufacturer’s protocol (10X Genomics, CG000183). The library was generated using the Chromium Single Cell 3’ Reagent Kits v3 (10X Genomics, PN-1000092) and Chromium i7 Multiplex Kit (10X Genomics, PN-120262) according to the manufacturer’s manual. Quality control for constructed library was performed by Agilent Bioanalyzer High Sensitivity DNA kit (Agilent Technologies, 5067-4626) for qualitative analysis. Quantification analysis was performed by Illumina Library Quantification Kit (KAPA Biosystems, KK4824). The library was sequenced on an Illumina HiSeq or NextSeq 2×150 paired-end kits using the following read length: 28 bp Read1 for cell barcode and UMI, 8 bp I7 index for sample index and 91 bp Read2 for transcript.

### Single cell ATAC sequencing

3-week-old and 8-week-old mice were euthanized and perfused with chilled 1x PBS via left ventricle. Kidneys (0.25 g) were harvested, minced and lysed in 5 mL lysis buffer for 15 min. 1-day-old mice were decapitated with surgical scissors, and both kidneys were harvested. Kidneys were minced and lysed in 2 mL lysis buffer for 15 min. Tissue lysis reaction was then blocked by adding 10 mL 1x PBS into each tube, and solution was passed through a 40 μm cell strainer. Cell debris and cytoplasmic contaminants were removed by Nuclei PURE Prep Nuclei Isolation Kit (Sigma, NUC-201) after centrifugation at 13,000 RPM for 45 min. Nuclei concentration was calculated with Countess AutoCounter (Invitrogen, C10227). Diluted nuclei suspension was loaded and incubated in transposition mix from Chromium Single Cell ATAC Library & Gel Bead Kit (10X Genomics, PN-1000110) by targeting 10,000 nuclei recovery. GEMs were then captured on the Chromium Chip E (10x Genomics, PN-1000082) in the Chromium Controller according to the manufacturer’s protocol (10X Genomics, CG000168). Libraries were generated using the Chromium Single Cell ATAC Library & Gel Bead Kit and Chromium i7 Multiplex Kit N (10X Genomics, PN-1000084) according to the manufacturer’s manual. Quality control for constructed library was perform by Agilent Bioanalyzer High Sensitivity DNA kit. The library was sequenced on an Illumina HiSeq 2×50 paired-end kits using the following read length: 50 bp Read1 for DNA fragments, 8 bp i7 index for sample index, 16 bp i5 index for cell barcodes and 50 bp Read2 for DNA fragments.

### Bulk ATAC sequencing

Bulk ATAC-seq was performed as described earlier ^63,64^. Briefly, 50,000 nuclei/sample were tagmented with Tn5 transposase (Illumina) in 50 μl reaction volume including Tween-20 (0.1%) (Sigma) and digitonin (0.01%) (Promega). The reaction was carried out at 37 °C for 30 min in a thermomixer at 1,000 RPM. After purification of DNA with Qiagen Minelute Reaction Cleanup kit (Qiagen), samples were subjected to library amplification (8-10 cycles). Libraries were purified with AmpureXP beads (Beckman Coulter) and their quality was assessed by Agilent High sensitivity DNA Bioanalysis chip (Agilent). Libraries were submitted to 150 bp PE sequencing.

### snATAC-seq data analysis

#### Data processing and quality control

Raw fastq files were aligned to the mm10 (GRCm38) reference genome and quantified using Cell Ranger ATAC (v. 1.1.0). We only kept valid barcodes with number of fragments ranging from 1,000 to 40,000 and mitochondria ratio less than 10%. One of the important indicators for ATAC-seq data quality is the fraction of peaks in promoter regions, so we did further filtration based on promoter ratio. We noticed the promoter ratio seemed to follow a binary distribution, with most of cells either having a promoter ratio around 5% (background) or more than 20% (valid cells) (**Figure S1d**). We therefore filtered out cells with a promoter ratio <20%. After this stringent quality control, we obtained 11,429 P0 single cells (5,993 in P0_batch_1 and 5,436 in P0_batch_2) and 16,887 adult single cells (7,129 in P56_batch_3, 6,397 in P56_batch_4, and 3,361 in P21_batch_5).

#### Preprocessing

Since snATAC-seq data are very sparse, previous methods either conducted peak calling or binarization before clustering. Here, we chose to do binarization instead of peak calling for two reasons: 1) Peak calling is time consuming; 2) Many peaks are cell type-specific, open chromatin regions in rare populations are more likely to be treated as background. After binarizing fragments into 5 kb bins and removing the fragments not matched to chromosomes or aligned to the mitochondria, we binarized the cell-bin matrix. In order to only keep bins that were informative for clustering, we removed the top 5% most accessible bins and bins overlapping with ENCODE blacklist. The 484,606 remaining bins were used as input for clustering.

#### Dimension reduction, batch effect correction and clustering

Clustering was conducted using snapATAC ^16^, a single-cell ATAC-seq algorithm scalable to large dataset. Previous benchmarking evaluation has shown that snapATAC was one of the best-performing methods for snATAC-seq clustering ^65^. Diffusion map was applied as a dimension reduction method using function *runDiffusionMaps*. To remove batch effect, we used Harmony ^17^, in which the low dimensional embeddings obtained from the diffusion map were used as input. Harmony iteratively pulled batch-specific centroid to cluster centroid until convergence to remove the variability across batches. After batch correction, a graph was constructed using k Nearest Neighbor (kNN) algorithm with k=15, which was then used as input for Louvain clustering. We used the first 20 dimensions for the Louvain algorithm. The number of dimensions was chosen using a method recommended by snapATAC, although we noticed that the clustering results were similar among a series of dimensions from 18 to 30.

#### Cell type annotation

We used a published list of marker genes ^9,19^ to annotate kidney cell types. In order to infer gene expression of each cell type, we built a cell-gene activity score matrix by integrating all fragments that overlapped with gene transcript. We used GENCODE Mouse release VM16 ^22^ as reference annotation.

#### Peak calling and visualization

Peak calling was conducted for each cell type separately using MACS2 ^18^. We aggregated all fragments obtained from the same cell types to build a pseudo-bulk ATAC data and conducted peak calling with parameters “--nomodel --keep-dup all --shift 100 --ext 200 --qval 1e-2-B -- SPMR --call-summits”. By specifying “--SPMR”, MACS2 generated “fragment pileup per million reads” pileup files, which were converted to bigwig format for visualization using UCSC bedGraphToBigWig tool.

We also visualized public chromatin ChIP-seq data and RNA-seq data obtained from ENCODE Encyclopedia with the following identifiers: ENCFF338WZP, ENCFF872MVE, ENCFF455HPY, ENCFF049LRQ, ENCFF179NTO, ENCFF071PID, ENCFF746MFH, ENCFF563LOO, ENCFF184AYF, ENCFF107NQP, ENCFF465THI, ENCFF769XWI, ENCFF591DAX. The Six2 ChIP-seq data were obtained from ^66^ and the WGBS data were obtained from ^67^.

#### Genomic elements stratification

Mouse mm10 genome annotation files were download from UCSC Table Browser (https://genome.ucsc.edu/cgi-bin/hgTables) using GENCODE VM23. TSS upstream 5 kb regions were included as promoter regions, but the results were similar when using 2 kb upstream regions as promoters. We then studied the number of overlapped regions between open chromatin regions identified from the snATAC-seq and bulk ATAC-seq dataset and genome annotations. Since one open chromatin region could overlap with multiple genomic elements, we defined an order of genomic elements as exon > 5’-UTR > 3’-UTR > intron > promoter > distal elements. To be more specific, if one peak overlapped with both exon and 5’-UTR, the algorithm would count it as an exon-region peak.

#### Identification of differentially accessible regions

Peaks identified in each cell type were combined to build a union peak set. Overlapping peaks were then merged to one peak using *reduce* function from the GenomicRanges package. This resulted in 300,755 peaks, which were used to build binarized cell-by-peak matrix. Differentially accessible peaks (DAPs) for each cell type were identified by pairwise peak comparison.

Specifically, for each peak, we conducted a Fisher’s exact test between a cell type and each of the other cell types. To address multiple testing problem, we used the Benjamini-Hochberg approach (BH correction) to correct p values. Peaks with corrected p values below significance level (0.05) in all pairwise tests were defined as DAPs. In total, we obtained 60,683 DAPs, which were used for motif enrichment analysis.

#### Motif enrichment analysis

Motif enrichment analysis was conducted using DAPs by HOMER v4.10.4 ^27^ with parameters background=“automatic” and scan.size=300. We noticed that *de novo* motif identification only generated few significant results, so we focused on known motifs for our following study. We used the significance level of 0.05 for BH corrected p value to determine the enriched results. The motif enrichment results are provided in **Supplemental Table 3**.

#### Peak-peak correlation analysis

Peak-peak correlation analysis was conducted using Cicero ^26^. In order to find developmental stage-specific peak-peak correlations, the analysis was conducted for P0 and adult separately. Cicero uses Graphic Lasso with distance penalty to assess the co-accessibility between different peaks. Cicero analysis was conducted using the *run_cicero* function with default parameters. A heuristic cutoff of 0.25 score of co-accessibility was used to determine the connections between two peaks.

#### snATAC-seq trajectory analysis

snATAC-seq trajectory was conducted using Cicero, which extended Monocle3 to the snATAC-seq analysis. We obtained the preprocessed P0 snATAC-seq cell-peak matrix from snapATAC as input for Cicero and conducted dimension reduction using Latent Semantic Indexing (LSI) and visualized using UMAP. Trajectory graph was built using the function *learn_graph*. Batch effect was not observed between the two P0 batches, and the trajectory graph was consistent with cell type assignment with clustering analysis (**Figures S3 a-b**).

In order to study how open chromatin changes are associated with the cell fate decision, we first binned the cells into 15 groups based on their pseudotime and cell type assignment. Next, we studied the DAPs between each group and its ancestral group using the same methods described above. The number of newly open and closed chromatins were reported using pie charts. The exact peak locations are provided in the **Supplemental Table 7**.

#### Genes and gene ontology terms associated with snATAC-seq trajectory

Based on the binned trajectory graphs and DAPs between each group and its ancestral group, we next used GREAT tool ^68^ to study the enrichment of associated genes and gene ontology (GO) terms along the trajectory. We used the newly open or closed peaks as test regions and all the peaks from peak-calling output as the background regions for the analysis. The output can be found in the **Supplemental Table 8 and 9**.

#### Predict cis-regulatory elements

We implemented two methods to study cis-regulatory elements in the snATAC-seq data. The first method was inspired by Zhu et al. ^23^, which was based on the observation that there was co-enrichment in the genome between the snATAC-seq cell type-specific peaks and scRNA-seq cell type-specific genes. This method links a gene with a peak if 1) they were both specific in the same cell type, 2) they were in *cis*, meaning that the peak is in ±100 kb region of the TSS of the corresponding gene, and 3) the peak did not directly overlap with the TSS of the gene. This method successfully inferred several known distal elements such as for *Six2* and *Slc6a18* (**Figure S2g**).

Alternatively, we assessed the co-accessibility of two peaks. We implemented Cicero ^26^, which aggregates similar cells to obtain a set of “meta-cells” and address the issue with sparsity in the snATAC-seq data. We used *run_cicero* function with default parameters to predict cis-regulatory elements (CREs). Although it is recommended to use 0.25 as a cutoff for co-accessibility score, we noticed that this resulted in a great amount of CREs, which could contain many false positives. Thus, we used a more stringent score of 0.4 for the cutoff and retained 232,380 and 206,701 CRE links in the P0 and adult data, respectively.

### Bulk ATAC sequencing analysis

Bulk ATAC-seq raw fastq files were processed using the end-to-end tool ENCODE ATAC-seq pipeline (**Software and Algorithms**). This tool provided a standard workflow for ATAC-seq data quality control, adaptor removal, alignment, and peak calling. To obtain high quality ATAC-seq peaks, peak calling results from two biological replicates were compared and only those peaks that were present in both replicates were kept, which were further used to compare with snATAC-seq peaks.

### Correlation of bulk and single nuclei ATAC sequencing data

snATAC-seq reads were aggregated to a pseudo-bulk data for the comparison purpose. To prevent the effect of sex chromosome and mitochondria chromosome, reads from chromosome X, Y and M were excluded from our analysis. We used multiBigwigSummary tool from deeptools ^69^ to study the correlation between different samples. Specifically, the whole genome was binned into equally sized (10 kb) windows, and the reads in each bin were aggregated, generating a bin-read count vector for each of the sample. The correlation of these vectors was computed as a measure of pairwise similarity between samples.

To compare the number of peaks in these two datasets, we used as input the narrowpeak files from the snATAC-seq and bulk ATAC-seq analysis. We filtered out bulk ATAC-seq peaks with q value > 0.01 to be consistent with the snATAC-seq setting. Since the snATAC peaks were called after merging different time points, we also took the union set of bulk ATAC-seq peaks from different time points. We then used *findoverlap* function in GenomicRanges package ^70^ to find and report overlapped peaks.

### Comparison between single nuclei ATAC sequencing data and single cell RNA sequencing data

In order to compare the cluster assignment between snATAC-seq data and scRNA-seq data, we obtained the average gene expression values and peak accessibility in each cluster for P0 and adult samples separately. We next transformed snATAC-seq data by summing up the reads within gene body and 2 kb upstream regions to build gene activity score matrix, as suggested in Seurat ^20^. Then, we normalized the data and computed the mean expression and mean gene activity scores in each cell type, and calculated z scores of each gene. Pearson’s correlation coefficient was then calculated among top 3,000 highly variable genes between snATAC-seq data and scRNA-seq data.

We found high concordance between these two datasets in terms of cell type assignment (**Figure S4**).

### Single cell RNA sequencing data analysis

#### Alignment and quality control

Raw fastq files were aligned to the mm10 (Ensembl GRCm38.93) reference genome and quantified using CellRanger v3.1.0. Seurat R package v3.0 was used for data quality control, preprocessing and dimensional reduction analysis. After gene-cell data matrix generation of both P0 and adult datasets, matrices were merged and poor-quality cells with <200 or >3,000 expressed genes and mitochondrial gene percentages >50 were excluded, leaving 25,138 P0 and 18,498 adult cells for further analytical processing, respectively (**Figures S1j-k**).

#### Pre-processing, batch effect correction and dimension reduction

Data were normalized by RPM following log transformation and 3,000 highly variable genes were selected for scaling and principal component analysis (PCA). Harmony R package v1.0 ^17^ was used to correct batch effects. The top 20 dimensions of Harmony embeddings were used for downstream uniform manifold approximation and projection (UMAP) visualization and clustering (**Figures S1l-m**).

#### Cell clustering, identification of marker genes and differentially expressed genes

Louvain algorithm with resolution 0.4 was used to cluster cells, which resulted in 18 distinct cell clusters. A gene was considered to be differentially expressed if it was detected in at least 25% of one group and with at least 0.25 log fold change between two groups and the significant level of BH-adjusted p value <0.05 in Wilcoxon rank sum test was used. We used a list of marker genes ^9,19^ to manually annotate cell types. 2 distal convoluted tubule clusters were merged based on the marker gene expression, resulting in a total of 17 clusters (**Figures S1i, n, o**).

#### scRNA-seq trajectory analysis

##### Monocle3

To construct single cell pseudotime trajectory and to identify genes whose expression changed as the cells underwent transition, Monocle3 v0.1.3 ^71^ was applied to P0 cells of the following Seurat cell clusters: nephron progenitors (NP), proliferating cells, stroma-like cells, podocytes, loop of Henle (LOH), early proximal tubule (PT), proximal tubule S1, proximal tubule S3 cells.

To show cell trajectories from both small (nephron progenitors) and large cell populations (proximal tubule), an equal number of 450 cells per cluster was randomly subsampled. Cells were re-clustered by Monocle3 using a resolution of 0.0005 with k-nearest neighbor (kNN) k=29. Highly variable genes along pseudotime were identified using differential *GeneTest* function and cells were ordered along pseudotime trajectory. NP cluster was defined as earliest principal node. In order to find genes differentially expressed along pseudotime, trajectories for podocytes, loop of Henle, and proximal tubule clusters were analyzed separately with the *fit_models* function of Monocle3. Genes with a q value <0.05 in the differential *GeneTest* analysis were kept. In an alternate approach, *graph_test* function of Monocle3 was used and trajectory-variable genes were collected into modules at a resolution of 0.01.

##### RNA velocity

To calculate RNA velocity, Python-based Velocyto command-line tool as well as Velocyto.R package were used as instructed ^30^. We used Velocyto to calculate the single-cell trajectory/directionality using spliced and unspliced reads. From loom files produced by the command-line tool, we subset the exact same cells that were previously selected randomly for Monocle trajectory analysis. This subset was loaded into R using the SeuratWrappers v0.1.0 package. RNA velocity was estimated using gene-relative model with k-nearest neighbor (kNN) cell pooling (k = 25). The parameter n was set at 200, when visualizing RNA velocity on the UMAP embedding.

#### Gene regulatory network inference

In order to identify TFs and characterize cell states, we employed *cis*-regulatory analysis using the R package SCENIC v1.1.2.2 ^28^, which infers the gene regulatory network based on co-expression and DNA motif analysis. The network activity is then analyzed in each cell to identify recurrent cellular states. In short, TFs were identified using GENIE3 and compiled into modules (regulons), which were subsequently subjected to *cis*-regulatory motif analysis using RcisTarget with two gene-motif rankings: 10 kb around the TSS and 500 bp upstream. Regulon activity in every cell was then scored using AUCell. Finally, binarized regulon activity was projected onto Monocle3-created UMAP trajectories.

#### Ligand-receptor interactions

To assess cellular crosstalk between different cell types, we used the CellPhoneDB repository to infer cell-cell communication networks from single cell transcriptome data ^40^. We used the Python package CellPhoneDB v2.1.2 together with the database v2.0.0 to predict cell type-specific ligand-receptor complexes as per the authors’ instructions. Only receptors and ligands expressed in more than 5% of the cells in the specific cluster were considered. 1,000 iterations were used for pairwise comparison between cell types and considered for further statistical analysis.

### Immunofluorescence staining

Mouse kidneys were fixed with 4% paraformaldehyde overnight, rinsed in PBS, and dehydrated for paraffin embedding. Antigen retrieval was performed using Tris-EDTA buffer pH 9.0 with a pressure cooker (PickCell Laboratories, Agoura Hills, CA) and antibody staining performed as described ^72^. Antibodies used were as follows: guinea pig FOXL1 (1:1,500) ^73^, mouse E-cadherin (1:250; BD Transducton 610182, Franklin Lakes, NJ). Cy2-, Cy3-, and Cy5-conjugated donkey secondary antibodies were purchased from Jackson ImmunoResearch Laboratories, Inc. Fluoresecence images were collected on a Keyence microscope.

## Material Table

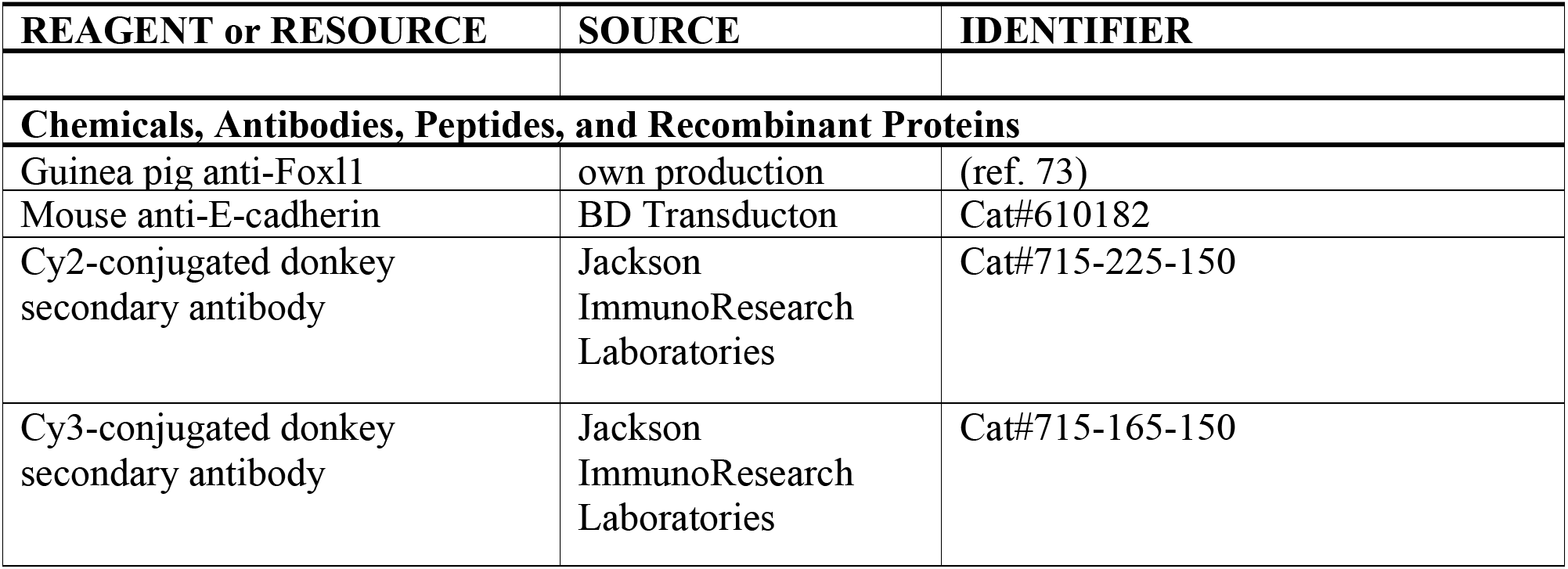
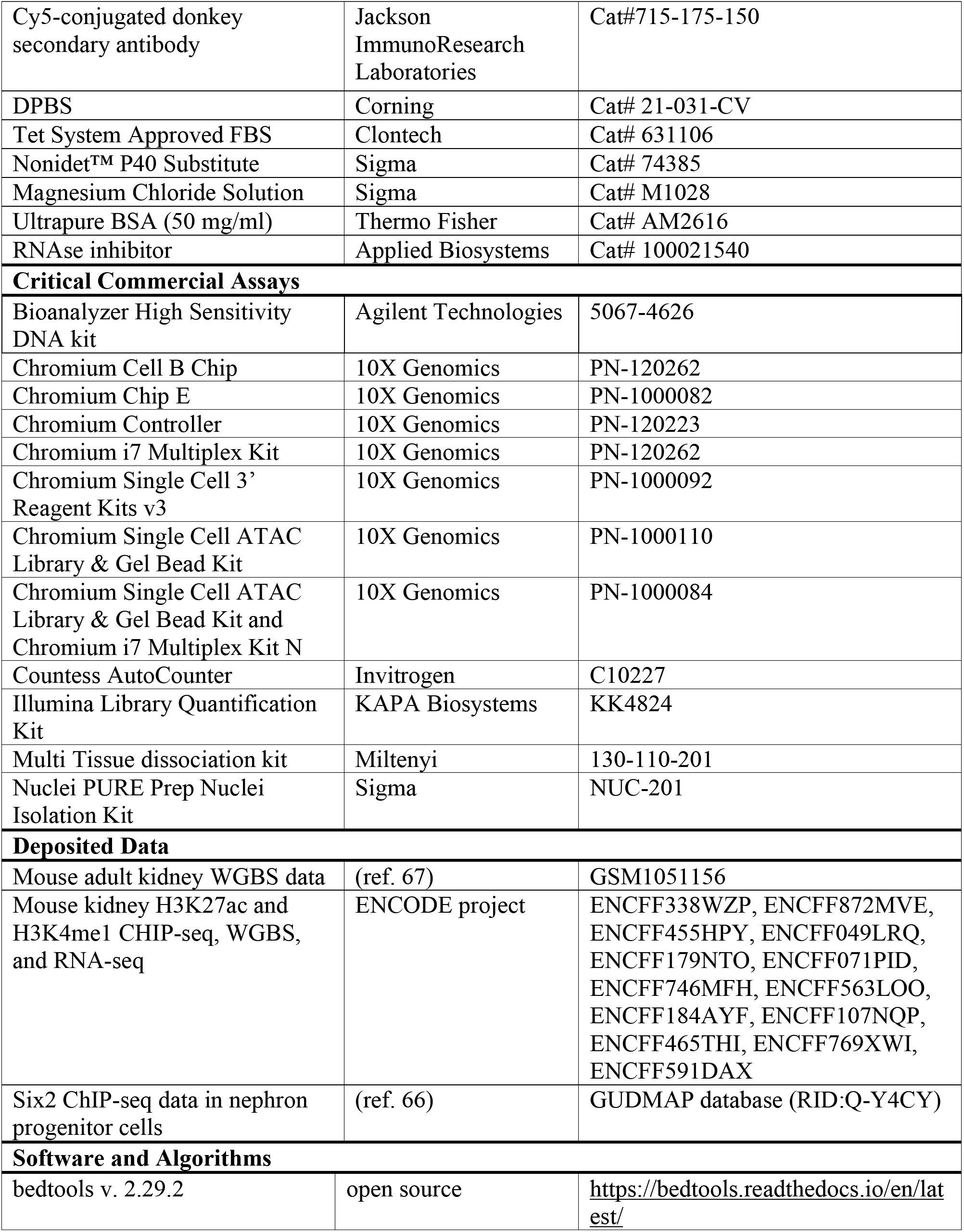
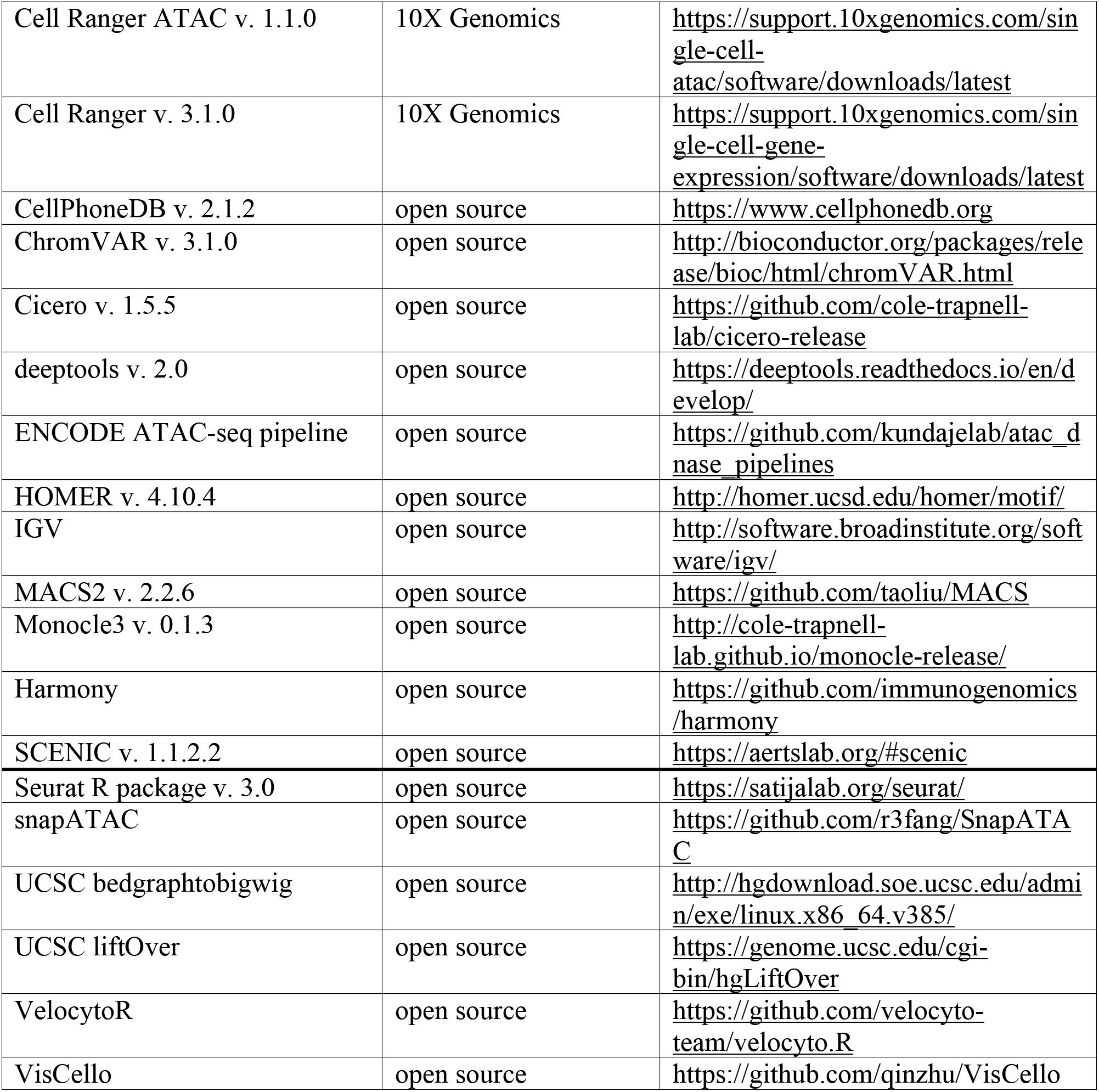

### Lead Contact and Data Availability

Raw data files and data matrix are being uploaded onto GEO and an accession number will be provided when it becomes available. The annotated and analyzed data can be viewed at susztaklab.com/developing_adult_kidney/snATAC/, susztaklab.com/developing_adult_kidney/scRNA/, and susztaklab.com/developing_adult_kidney/igv/. Further information and requests for resources and reagents should be directed to and will be fulfilled by the lead contact: Katalin Susztak.

## Supplemental Information

### Supplemental Tables

**Supplemental Table 1.** Cell type marker genes derived from scRNA-seq analysis.

**Supplemental Table 2.** Cell type-specific open chromatin derived from snATAC-seq analysis.

**Supplemental Table 3.** Cell type-specific motif enrichment.

**Supplemental Table 4.** Regulons and respective target genes inferred by SCENIC.

**Supplemental Table 5.** Binarized regulon activities in each cell type inferred by SCENIC.

**Supplemental Table 6.** Differentially expressed genes along pseudotime in distinct lineages in scRNA-seq data.

**Supplemental Table 7.** Differentially accessible peaks along pseudotime in distinct lineages in snATAC-seq data.

**Supplemental Table 8**. Nearest genes of differentially accessible peaks along pseudotime in distinct lineages in snATAC-seq data.

**Supplemental Table 9**. GO enrichment of differentially accessible peaks along pseudotime in distinct lineages inferred by GREAT analysis.

**Supplemental Table 10.** ChromVAR cell-TF enrichment score matrix.

**Supplemental Table 11.** Nearest genes of differentially accessible peaks at bifurcation events along pseudotime in distinct lineages inferred by GREAT analysis.

**Supplemental Table 12.** Proportion of cells in each cell type with accessible chromatin overlapped with kidney disease associated SNPs.

### Supplemental Figures

**Figure S1. Quality control and data processing methods for snATAC-seq and scRNA-seq data analysis.**

(A) Insert size distribution of the 5 snATAC-seq samples showing periodic patterns.

(B) Transcription start sites (TSS) signal enrichment of the 5 snATAC-seq samples.

(C) Spearman correlation between snATAC-seq datasets and bulk ATAC-seq of binned genomic regions.

(D) Distribution of number of unique molecular identifiers (UMIs, x axis) and promoter ratio (y axis) in 5 samples shown by dot plot.

(E) Violin plots representing the number of accessible peaks across different clusters in the snATAC-seq dataset indicating similar distributions.

(F) UMAP representation of the snATAC-seq dataset colored by batches.

(G) Stacked bar graphs representing absolute numbers and percentages of identified cell types across snATAC-seq batches.

(H) Genome browser view of cell type-specific peaks at the TSS of marker genes for 13 cell types in the snATAC-seq dataset.

(I) From left to right: Stacked bar graphs showing the percentage of different cell types in the P0 and adult scRNA-seq datasets, tables showing the number of cells in each cell type (nCells) and corresponding percentage. NP, nephron progenitor; Podo, podocyte; PT, proximal tubule; S1, segment 1; S3, segment 3; LOH, loop of Henle; DCT, distal convoluted tubule cells; PC, collecting duct principal cells; IC, collecting duct intercalated cells; Endo, endothelial cells; Macro, macrophages; Neutro, neutrophils.

(J) UMAP representation of scRNA-seq data colored by the mitochondrial gene ratio (Mt %).

(K) Violin plots showing number of informative genes per single cell and unique molecular identifiers (UMIs) per single cell. Blue denotes adult kidney, orange denotes P0 kidney.

(L, M) Principal component (PC) representation of combined adult and P0 scRNA-seq dataset (left panel) and violin plots of corresponding embeddings values (right panel) before (L) and after (M) batch correction using Harmony.

(N) Dot plot of cell type-specific marker genes. Dot size denotes percentage of cells expressing the marker. Color scale represents average expression, orange denotes P0, blue denotes adult kidney.

(O) Feature plots of representative marker genes projected on UMAP dimension.

(P) Correlation between snATAC-seq gene activity scores and gene expression values in adult data, which is complementary to **Figure 1g**.

(Q) We provide the processed chromatin accessibility dataset via a searchable, interactive website (susztaklab.com/developing_adult_kidney/igv/). Ace2 was used as an example, and we show proximal tubule-specific enrichment of peaks at transcription start sites of the *Ace2* (Angiotensin-converting enzyme 2) gene (red boxes).

**Figure S2. Characterization of the cell type-specific regulatory landscape**

(A) Bar graph representing the number of accessible peaks in distal elements, promoters, introns, 5’-UTR, 3’UTR and exons, as distributed across samples of snATAC-seq data and bulk ATAC-seq data.

(B) Overlap of scATAC-seq differentially accessible peaks among cell types with H3K27Ac ChIP-seq data.

(C) Number of shared and unique peaks among snATAC-seq cell types. Cell types include nephron progenitors and cells differentiated from nephron progenitors.

(D) Genome browser view of Umod as an example for distal open chromatin region and its target promoter region.

(E) Distribution of different open chromatin elements in snATAC-seq cell types.

(F) Distribution of different open chromatin elements among differentially accessible peaks (DAPs) in snATAC-seq cell types.

(G) Genome browser representations of single cell open chromatin data for individual cell types at chromosomal loci around Six2 and Slc6a18, along with their known distal elements (red boxes). Corresponding chromosomal interaction of open chromatin regions, as inferred by Cicero (**Methods**), is depicted at the top.

(H) Genome browser views of representative marker genes demonstrating cell type-specific chromatin accessibility for proximal tubule (*Hnf4a* and *Hmgb3*), several tubular segments (*Hnf1b*), loop of Henle and distal convoluted tubule (*Esrrb* and *Ppargc1a*) as well as nephron progenitors and podocytes (Wt1).

(I) UMAP depiction of regulon activity (“on-blue”, “off-grey”) and RNA expression (red scale) of exemplary regulons of proximal tubule (*Hnf1a*), nephron progenitors (*Six2*), loop of Henle (*Ppargc1a*), proliferating cells (*Hmgb3*) and podocytes (*Mafb*), respectively. Exemplary target gene expression for the respective TF is shown in purple scale.

(J, K) Bar graphs depicting the absolute number of cell type-specific TFs reported by HOMER (J) and SCENIC (K) cis-regulatory analyses, respectively, as well as the number of TFs among DEGs from RNA expression data alone.

**Figure S3. snATAC-seq and scRNA-seq cell differentiation trajectories.**

(A) UMAP representation of snATAC-seq trajectory lineages of podocytes, proximal tubule and loop of Henle cells from nephron progenitors colored by 2 P0 batches.

(B) UMAP representation of snATAC-seq trajectory lineages of podocytes, proximal tubule and loop of Henle cells from nephron progenitors colored by original cell type assignment as in **Figure 1b**.

(C) UMAP representation of scRNA-seq trajectory lineages of podocytes, proximal tubule and loop of Henle cells from nephron progenitors colored by original cell type assignment as in **Figure 1b**.

(D) UMAP representation of RNA velocity of scRNA-seq trajectory inferred by VelocytoR, colored by original cell type assignment. Each dot is one cell and each arrow represents the time derivative of the gene expression state.

(E) UMAP representation of snATAC-scRNA integration results colored by cell type assignment.

(F) UMAP representation of snATAC-scRNA integration results colored by technologies (snATAC=red, scRNA=grey). Podo: podocytes, PT: proximal tubule, LOH: loop of Henle, DCT: distal convoluted tubule, NP: nephron progenitors, IM: intermediate stage cells.

(G) Dot plot showing snATAC-scRNA integration cell type assignment confusion matrix. Each column represents the original cell type assignment of snATAC-seq data, and each row represents the predicted cell type assignment by the integration analysis scRNA-seq data. Each dot represents the number of cells that were matched in the integrated data.

(H) Heatmap of chromVAR enrichment results. The original data matrix is given in **Supplemental Table 10**.

(I) Pseudotime-dependent chromatin accessibility and gene expression changes along the proximal tubule (red), podocytes (green) and loop of Henle (blue) cell lineages. The first column represents the dynamics of chromVAR TF enrichment score, the second column represents the dynamics of TF gene expression values, and the third and fourth column represent the dynamics of SCENIC-reported target gene expression values.

**Figure S4. Chromatin dynamics of nephron progenitor differentiation.**

(A) Di-graph representing cell type and lineage divergence, as derived from Cicero trajectory inference. Nephron progenitors (NP), podocytes (Podo), intermediate stage (IM), proximal tubule (PT), loop of Henle (LOH) and distal convoluted tubule (DCT) are connected with their developmental precursor stages and ordered by ascending numbering. Pie charts represent differentially assessible peaks (DAPs) between two stages, where the size of pie charts is proportional to the number of DAPs, orange color represents the number of open peaks, grey color the number of closed peaks. Bar graphs depict gene ontology (GO) term analysis of genes nearby DAPs derived from GREAT analysis (full list in **Supplemental Table 9**).

(B) Immunofluorescence staining of fetal mouse kidney. Upper panel and insert denote E13.5 stage, lower panel denotes P6 mouse. Blue staining represents nuclei (DAPI), green staining represents tubular epithelium (E-Cadherin) and red staining represents progenitor cells (FOXL1) along a developmental trajectory from early progenitor stage (asterisk) over comma-shaped (+) and S shaped bodies (cross) towards podocytes within primitive glomeruli (#).

(C) Pseudotime-dependent chromatin accessibility and gene expression changes along the proximal tubule (red), podocytes (green) and loop of Henle (LOH, blue) cell lineages for important bifurcation TFs in the podocyte (*Foxl1*) and distal tubule (*Tfap2b*) lineage.

(D) Bar graphs denote the percentage of cells with accessible chromatin of several *Six2* promoters and enhancers (gene loci numbered 1-3) as well as putative *Foxl1* enhancers (gene loci numbered 4-7) along pseudotime. Exact gene loci of enhancers and promoters are given above each respective graph. Changes along pseudotime are depicted for 3 lineages from nephron progenitors (NP) to podocytes, proximal tubule (PT) and loop of Henle (LOH) cells, respectively. The right upper subpanel depicts the genome browser overview of chromatin accessibility for the NP and therefore corresponds to the first bar in graphs on the left. The right lower subpanel depicts zoom-in versions of the 7 loci for all 3 lineages.

**Figure S5. Accessibility of peaks overlap with kidney disease SNPs.**

Heatmap showing the proportion of cells in each cell type with accessibility of SNP-overlapped peaks. The SNP IDs as well as nearest genes are provided in **Supplemental Table 12**.

**Figure S6. Single cell level chromatin accessibility highlighted human kidney GWAS target genes and cell types.**

Open chromatin and co-accessibility view at alternative scales to those shown in **Figure 6**.

(A, C, E) From top to bottom: Cicero-inferred co-accessibility of open chromatin regions in mouse orthologues of human *Shroom3*, *Dab2* and *Uncx* loci; Gene browser view of the single nucleotide polymorphisms within the regions; gene browser view of chromatin accessibility for nephron progenitors (NP), collecting duct intercalated cells (IC), collecting duct principal cell types (PC), proximal tubules segment 1 and 3 (PT S1 and PT S3), loop of Henle (LOH), distal convoluted tubule (DCT), stromal cells (stroma), podocytes (Podo), endothelial cells (Endo) and immune cells (Immune). Right subpanel shows violin plots of scRNA-seq gene expression in P0 (orange) and adult (blue) kidneys.

(B, D, F) Whole kidney H3K27ac, H3K4me1 and Six2 ChIP-seq, whole genome bisulfate sequencing (WGBS) and RNA-seq data in E15.5, P0 and adult kidney samples.

