## Supplemental Figures for "Single cell resolution regulatory landscape of the mouse kidney highlights cellular differentiation programs and renal disease targets"

Figure S1

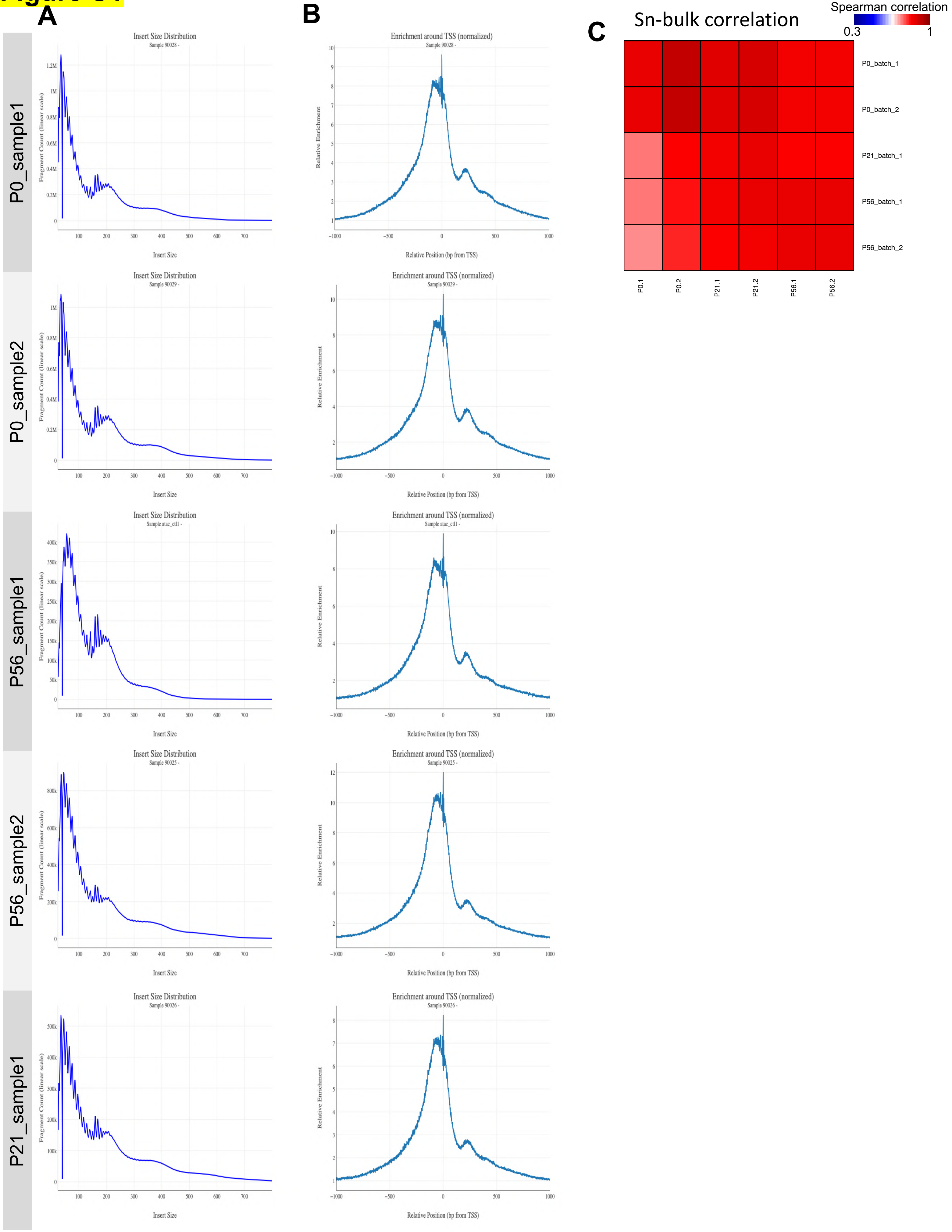

A
B
C
P0_sample1
P0_sample2
P56_sample1
P56_sample2
P21_sample1
Insert Size Distribution
Sample 90028 -
Sample 90029 -
Sample atac_ctl1 -
Sample 90025 -
Sample 90026 -
Fragment Count (linear scale)
Insert Size
Enrichment around TSS (normalized)
Relative Enrichment
Relative Position (bp from TSS)
Sn-bulk correlation
Spearman correlation
0.3
1
P0_batch_1
P0_batch_2
P21_batch_1
P56_batch_1
P56_batch_2
P0.1
P0.2
P21.1
P21.2
P56.1
P56.2

**Figure S1**

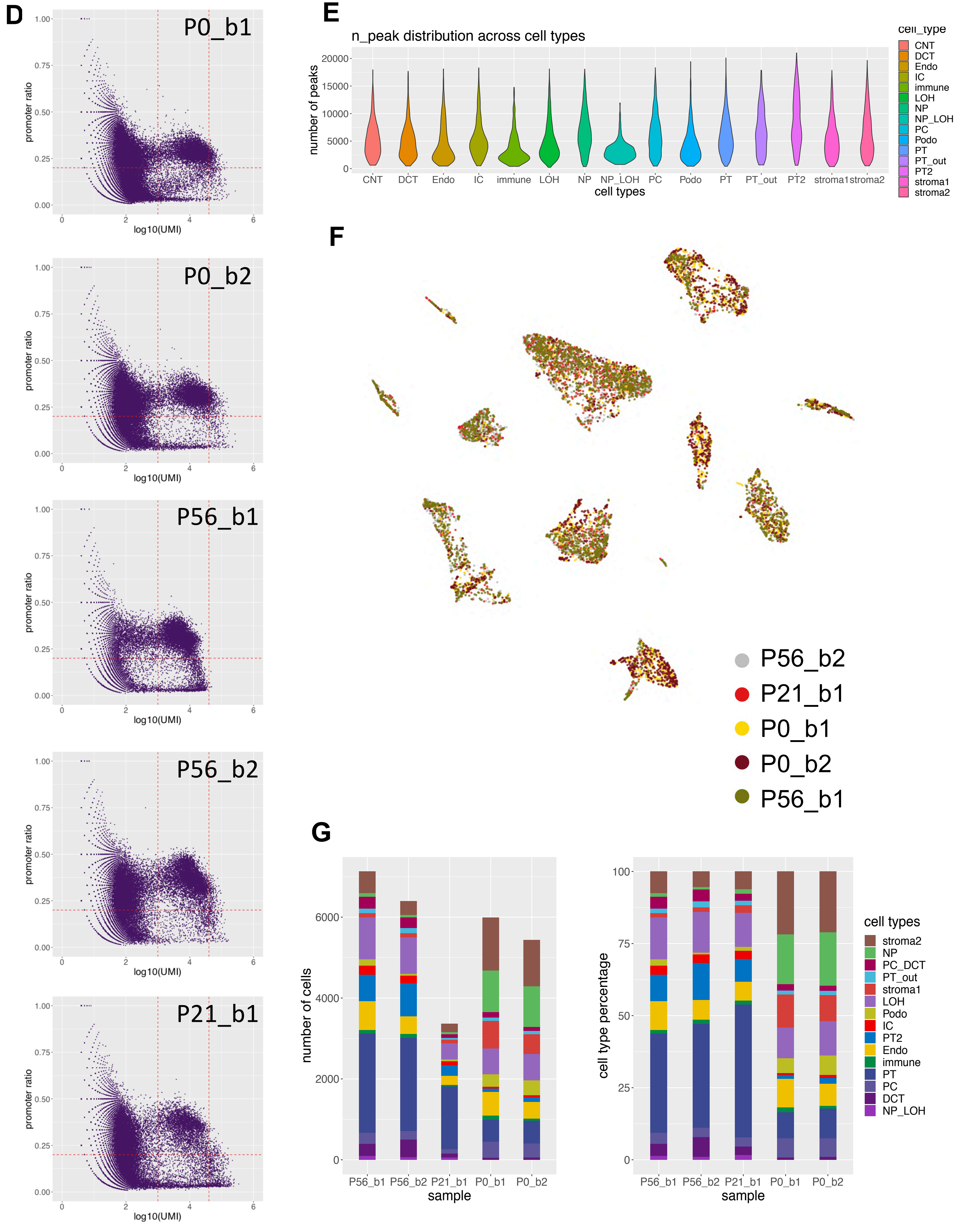

D
P0_b1
P0_b2
P56_b1
P56_b2
P21_b1
promoter ratio
log10(UMI)
E
n_peak distribution across cell types
number of peaks
cell types
CNT
DCT
Endo
IC
immune
LOH
NP
NP_LOH
PC
Podo
PT
PT_out
PT2
stroma1
stroma2
cell_type
F
P56_b2
P21_b1
P0_b1
P0_b2
P56_b1
G
number of cells
cell type percentage
sample
cell types
stroma2
NP
PC_DCT
PT_out
stroma1
LOH
Podo
IC
PT2
Endo
immune
PT
PC
DCT
NP_LOH

Figure S1

H

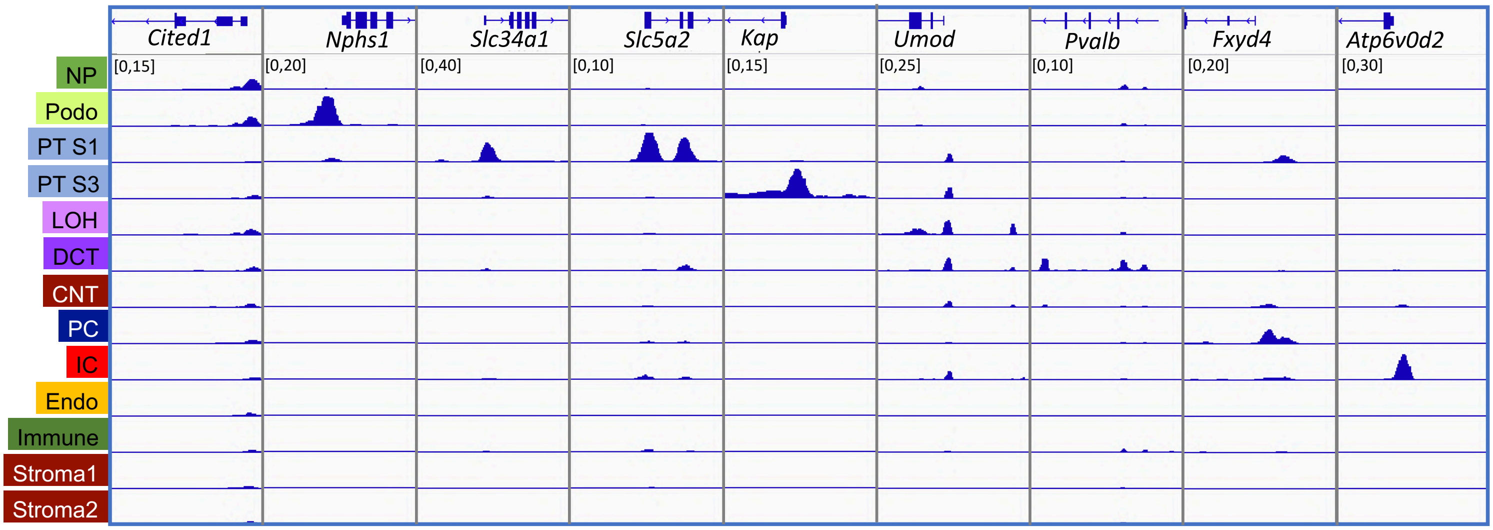

Cited1
Nphs1
Slc34a1
Slc5a2
Kap
Umod
Pvalb
Fxyd4
Atp6v0d2
[0,15]
[0,20]
[0,40]
[0,10]
[0,15]
[0,25]
[0,10]
[0,20]
[0,30]
NP
Podo
PT S1
PT S3
LOH
DCT
CNT
PC
IC
Endo
Immune
Stroma1
Stroma2

### Figure S1

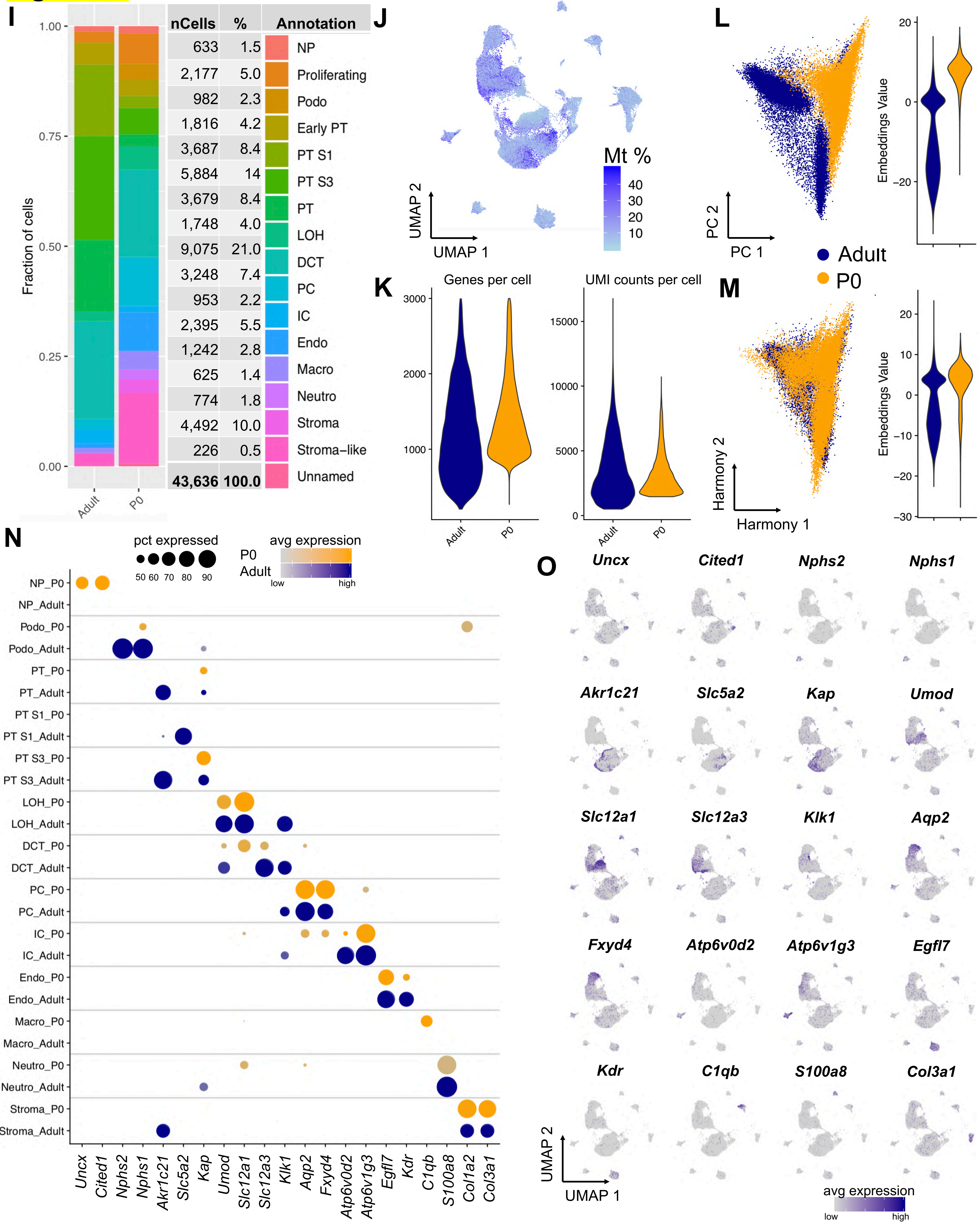
I
Fraction of cells
1.00
0.75
0.50
0.25
0.00
Adult
P0
nCells | % | Annotation
633 | 1.5 | NP
2,177 | 5.0 | Proliferating
982 | 2.3 | Podo
1,816 | 4.2 | Early PT
3,687 | 8.4 | PT S1
5,884 | 14 | PT S3
3,679 | 8.4 | PT
1,748 | 4.0 | LOH
9,075 | 21.0 | DCT
3,248 | 7.4 | PC
953 | 2.2 | IC
2,395 | 5.5 | Endo
1,242 | 2.8 | Macro
625 | 1.4 | Neutro
774 | 1.8 | Stroma
4,492 | 10.0 | Stroma–like
226 | 0.5 | Unnamed
43,636 | 100.0
J
Mt %
40
30
20
10
UMAP 2
UMAP 1
K
Genes per cell
UMI counts per cell
3000
2000
1000
15000
10000
5000
0
Adult
P0
L
PC 2
PC 1
Embeddings Value
20
0
−20
Adult
P0
M
Harmony 2
Harmony 1
Embeddings Value
20
10
0
−10
−20
−30
N
pct expressed
50 60 70 80 90
avg expression
P0
Adult
low
high
NP_P0
NP_Adult
Podo_P0
Podo_Adult
PT_P0
PT_Adult
PT S1_P0
PT S1_Adult
PT S3_P0
PT S3_Adult
LOH_P0
LOH_Adult
DCT_P0
DCT_Adult
PC_P0
PC_Adult
IC_P0
IC_Adult
Endo_P0
Endo_Adult
Macro_P0
Macro_Adult
Neutro_P0
Neutro_Adult
Stroma_P0
Stroma_Adult
Uncx Cited1 Nphs2 Nphs1 Akr1c21 Slc5a2 Kap Umod Slc12a1 Slc12a3 Klk1 Aqp2 Fxyd4 Atp6v0d2 Atp6v1g3 Egfl7 Kdr C1qb S100a8 Col1a2 Col3a1
O
Uncx
Cited1
Nphs2
Nphs1
Akr1c21
Slc5a2
Kap
Umod
Slc12a1
Slc12a3
Klk1
Aqp2
Fxyd4
Atp6v0d2
Atp6v1g3
Egfl7
Kdr
C1qb
S100a8
Col3a1
UMAP 2
UMAP 1
avg expression
low
high

Figure S1

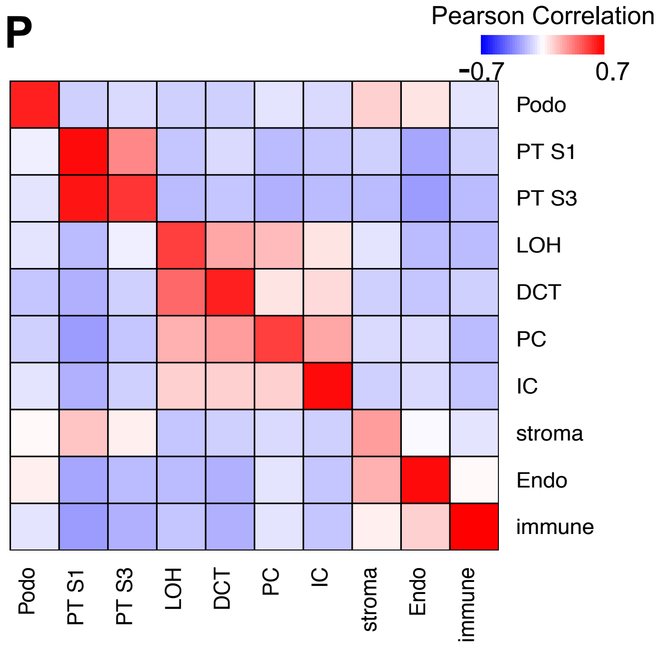
P
Pearson Correlation
-0.7
0.7
Podo
PT S1
PT S3
LOH
DCT
PC
IC
stroma
Endo
immune

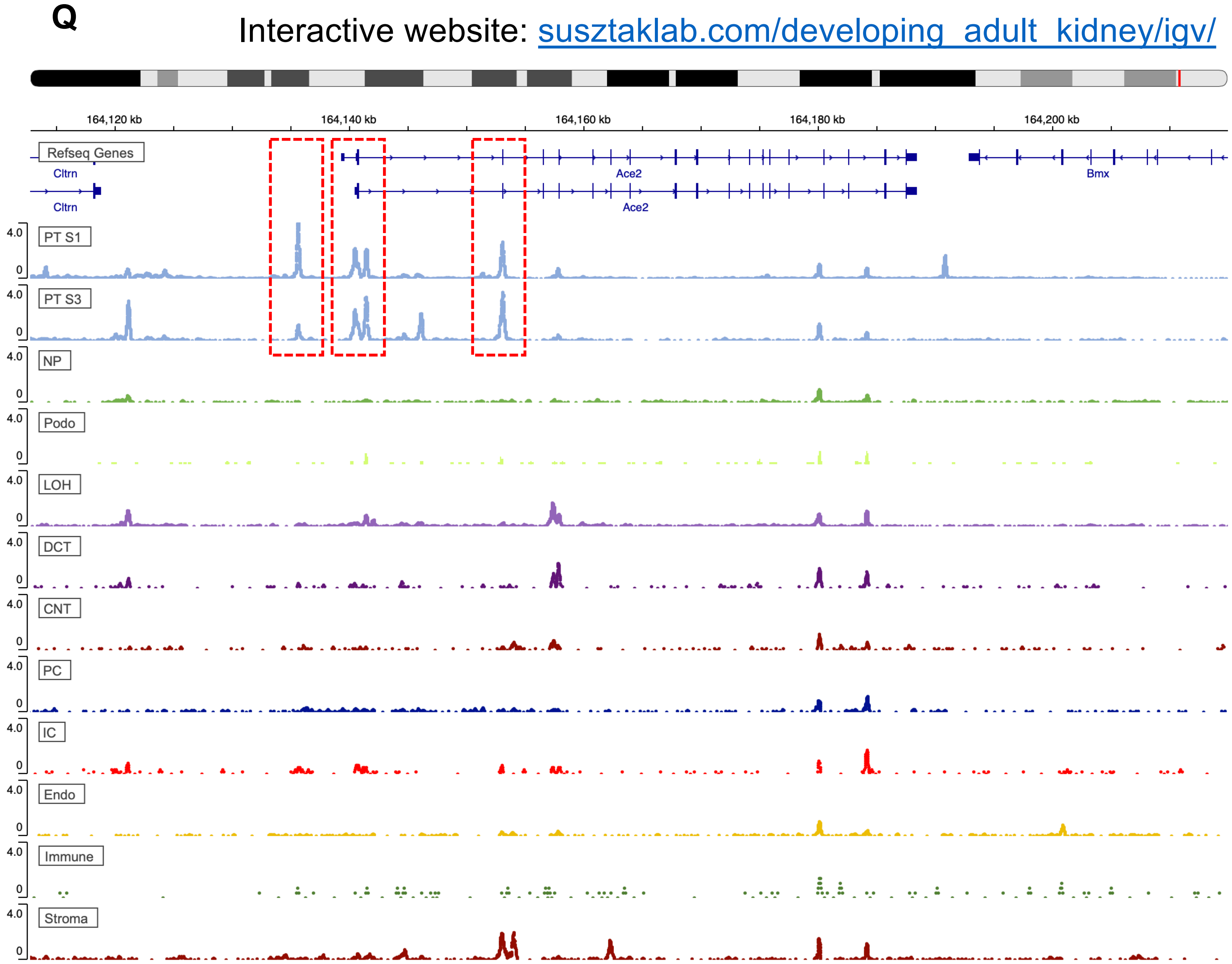
Q
Interactive website: susztaklab.com/developing_adult_kidney/igv/
164,120 kb
164,140 kb
164,160 kb
164,180 kb
164,200 kb
Refseq Genes
Cltrn
Ace2
Bmx
4.0
0
PT S1
PT S3
NP
Podo
LOH
DCT
CNT
PC
IC
Endo
Immune
Stroma

### Figure S2

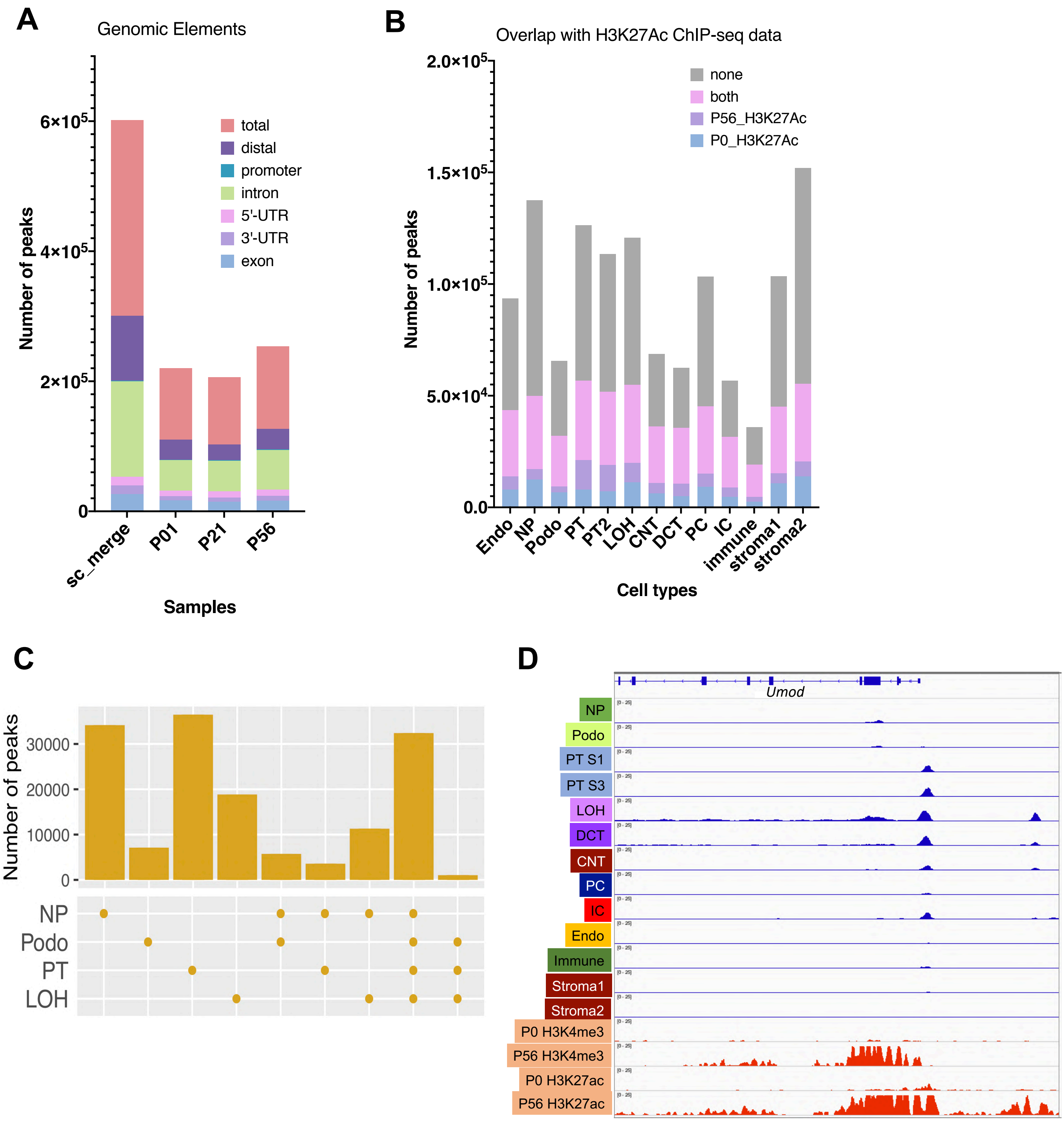
A
Genomic Elements
Number of peaks
6×10^5
4×10^5
2×10^5
0
total
distal
promoter
intron
5'-UTR
3'-UTR
exon
sc_merge
P01
P21
P56
Samples
B
Overlap with H3K27Ac ChIP-seq data
2.0×10^5
1.5×10^5
1.0×10^5
5.0×10^4
0.0
none
both
P56_H3K27Ac
P0_H3K27Ac
Number of peaks
Endo
NP
Podo
PT
PT2
LOH
CNT
DCT
PC
IC
immune
stroma1
stroma2
Cell types
C
Number of peaks
30000
20000
10000
0
NP
Podo
PT
LOH
D
Umod
NP
Podo
PT S1
PT S3
LOH
DCT
CNT
PC
IC
Endo
Immune
Stroma1
Stroma2
P0 H3K4me3
P56 H3K4me3
P0 H3K27ac
P56 H3K27ac

**Figure S2**

E

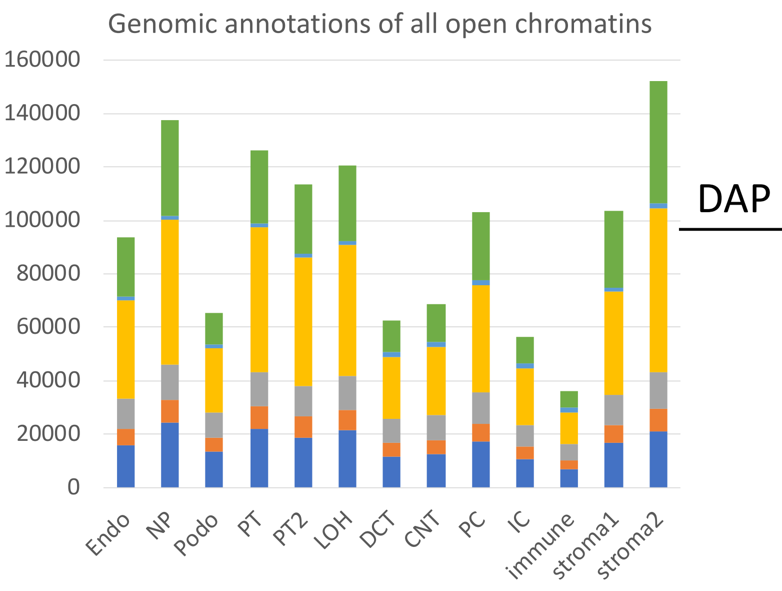


F

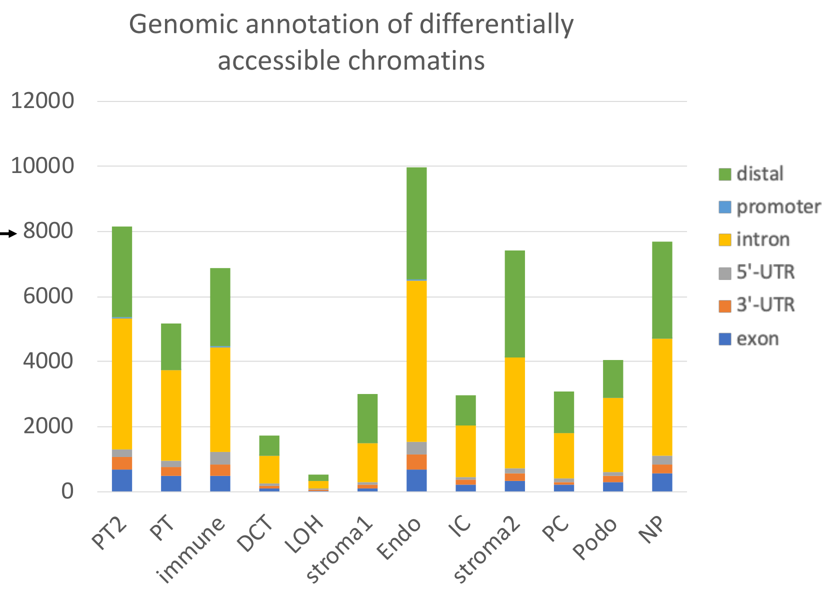


G

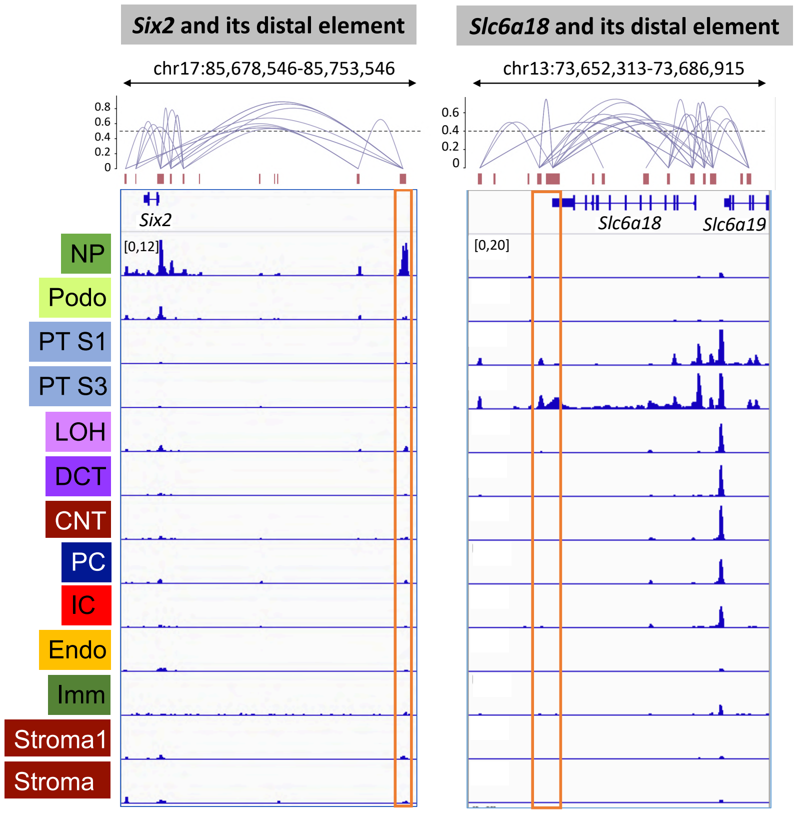

### Figure S2

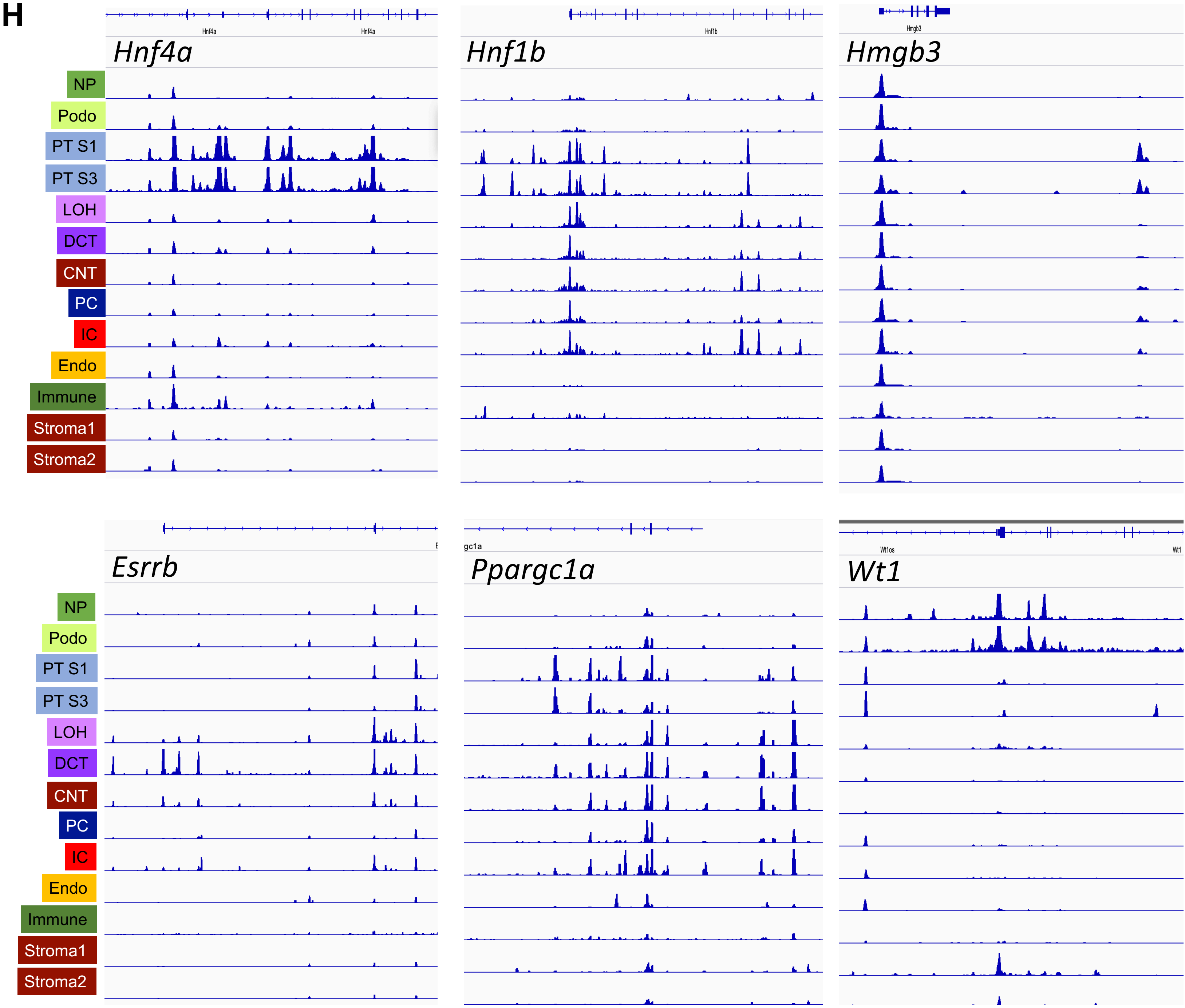
H
Hnf4a
Hnf1b
Hmgb3
Esrrb
Ppargc1a
Wt1
NP
Podo
PT S1
PT S3
LOH
DCT
CNT
PC
IC
Endo
Immune
Stroma1
Stroma2

### Figure S2

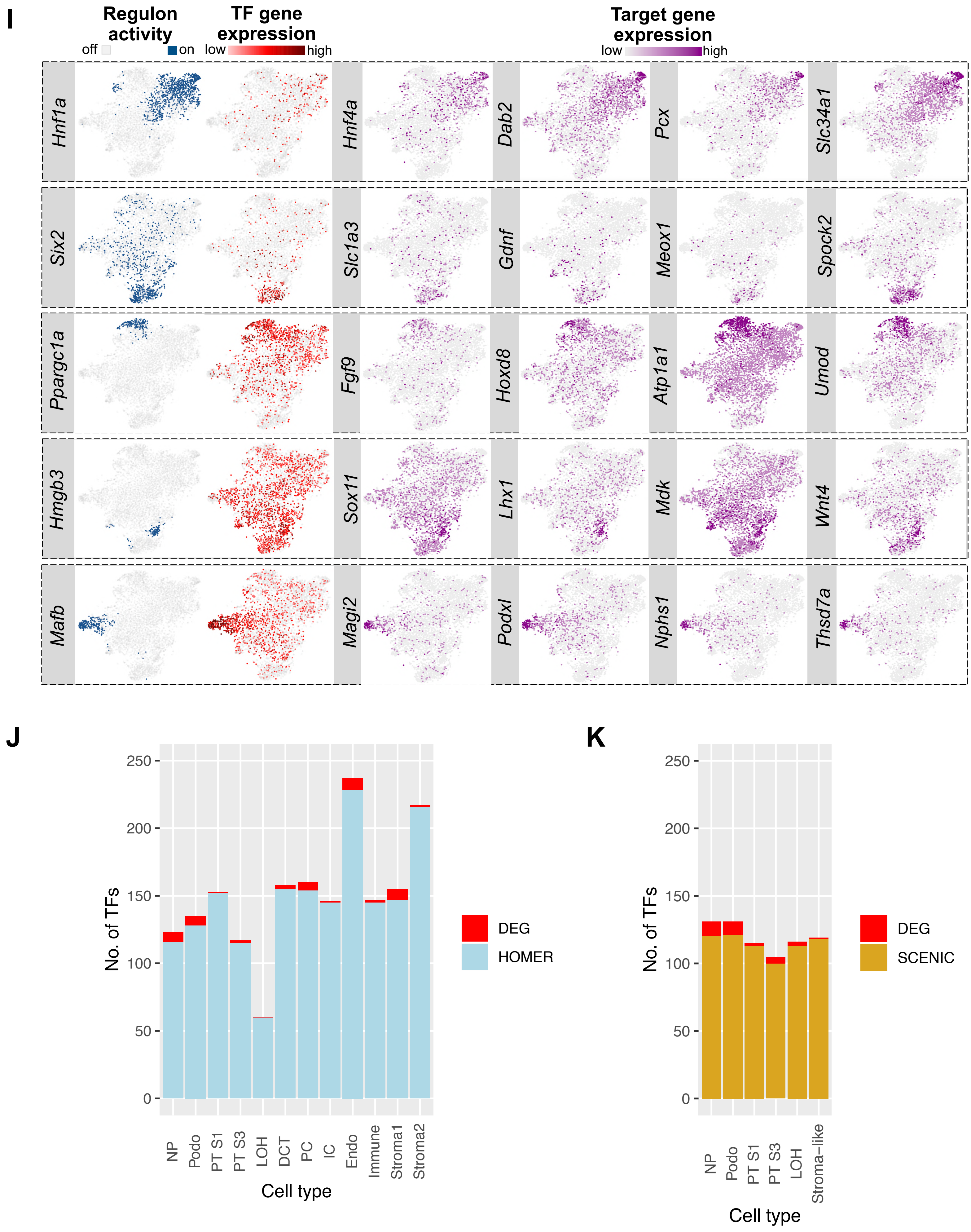
I
Regulon activity
off
on
TF gene expression
low
high
Target gene expression
low
high
Hnf1a
Hnf4a
Dab2
Pcx
Slc34a1
Six2
Slc1a3
Gdnf
Meox1
Spock2
Ppargc1a
Fgf9
Hoxd8
Atp1a1
Umod
Hmgb3
Sox11
Lhx1
Mdk
Wnt4
Mafb
Magi2
Podxl
Nphs1
Thsd7a
J
No. of TFs
0
50
100
150
200
250
NP
Podo
PT S1
PT S3
LOH
DCT
PC
IC
Endo
Immune
Stroma1
Stroma2
Cell type
DEG
HOMER
K
No. of TFs
0
50
100
150
200
250
NP
Podo
PT S1
PT S3
LOH
Stroma−like
Cell type
DEG
SCENIC

Figure S3

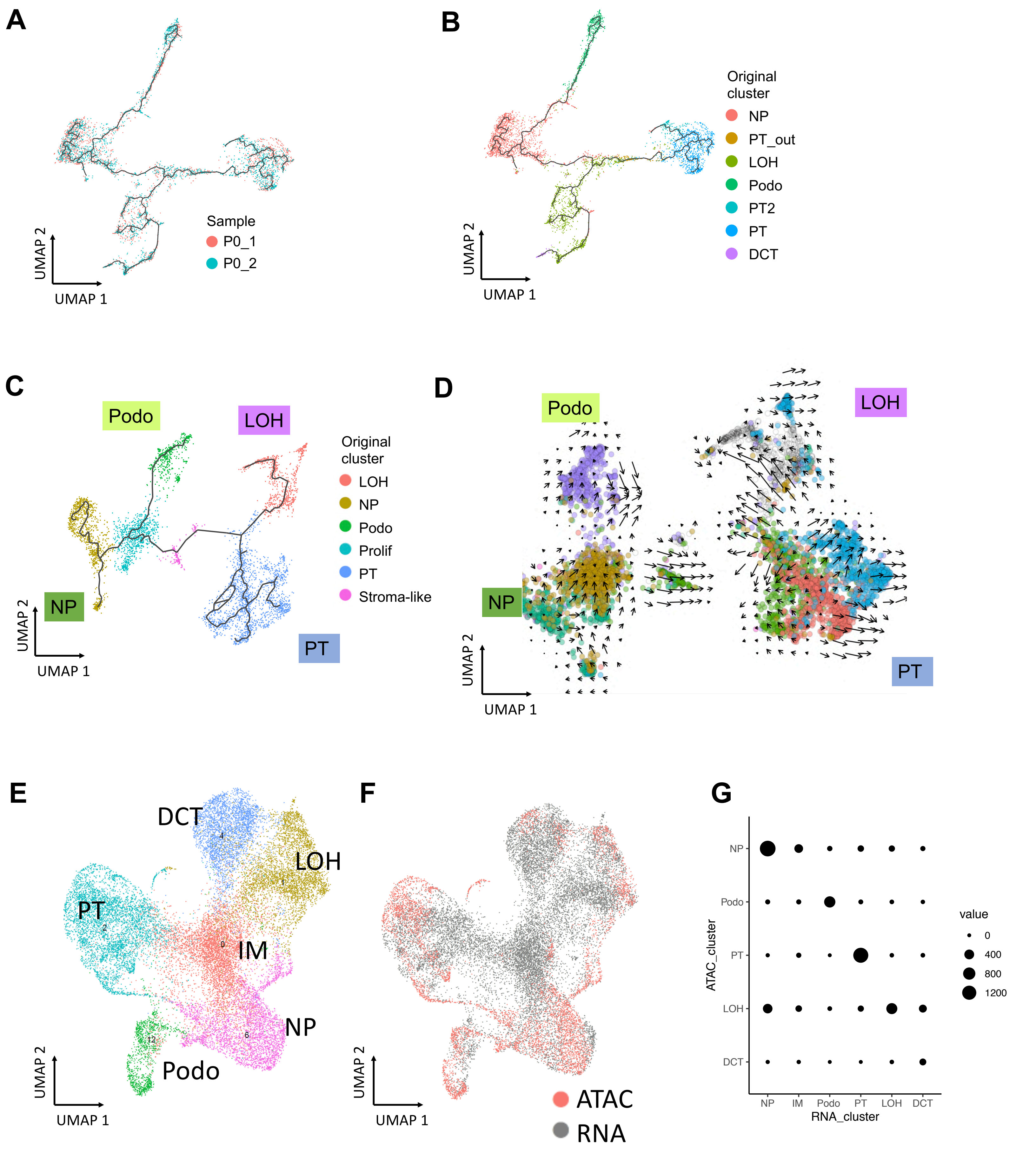
A
Sample
P0_1
P0_2
UMAP 2
UMAP 1
B
Original cluster
NP
PT_out
LOH
Podo
PT2
PT
DCT
UMAP 2
UMAP 1
C
Podo
LOH
NP
PT
Original cluster
LOH
NP
Podo
Prolif
PT
Stroma-like
UMAP 2
UMAP 1
D
Podo
LOH
NP
PT
UMAP 2
UMAP 1
E
DCT
LOH
PT
IM
NP
Podo
UMAP 2
UMAP 1
F
ATAC
RNA
UMAP 2
UMAP 1
G
NP
Podo
PT
LOH
DCT
ATAC_cluster
NP
IM
Podo
PT
LOH
DCT
RNA_cluster
value
0
400
800
1200

### Figure S3

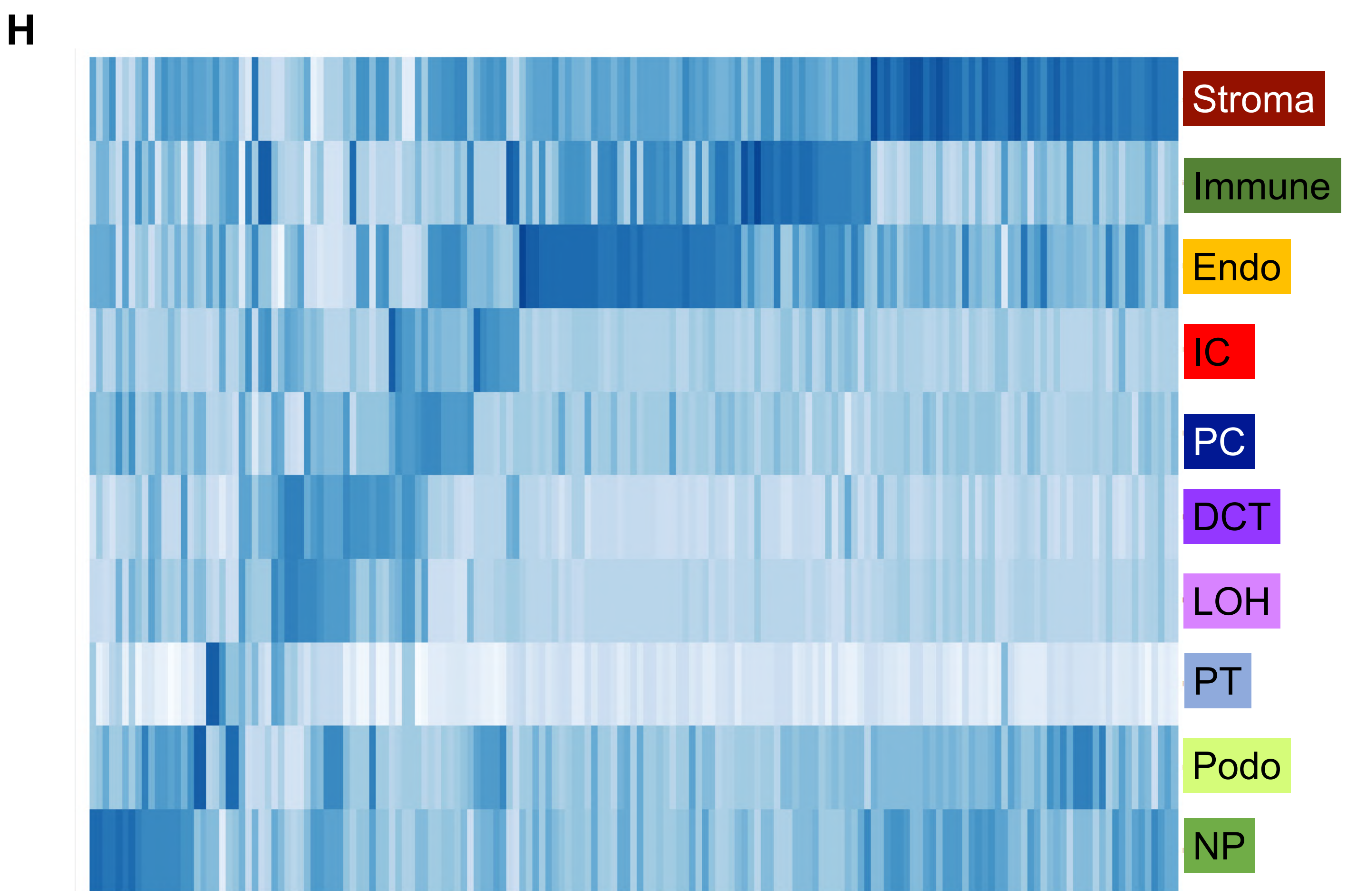
H
Stroma
Immune
Endo
IC
PC
DCT
LOH
PT
Podo
NP

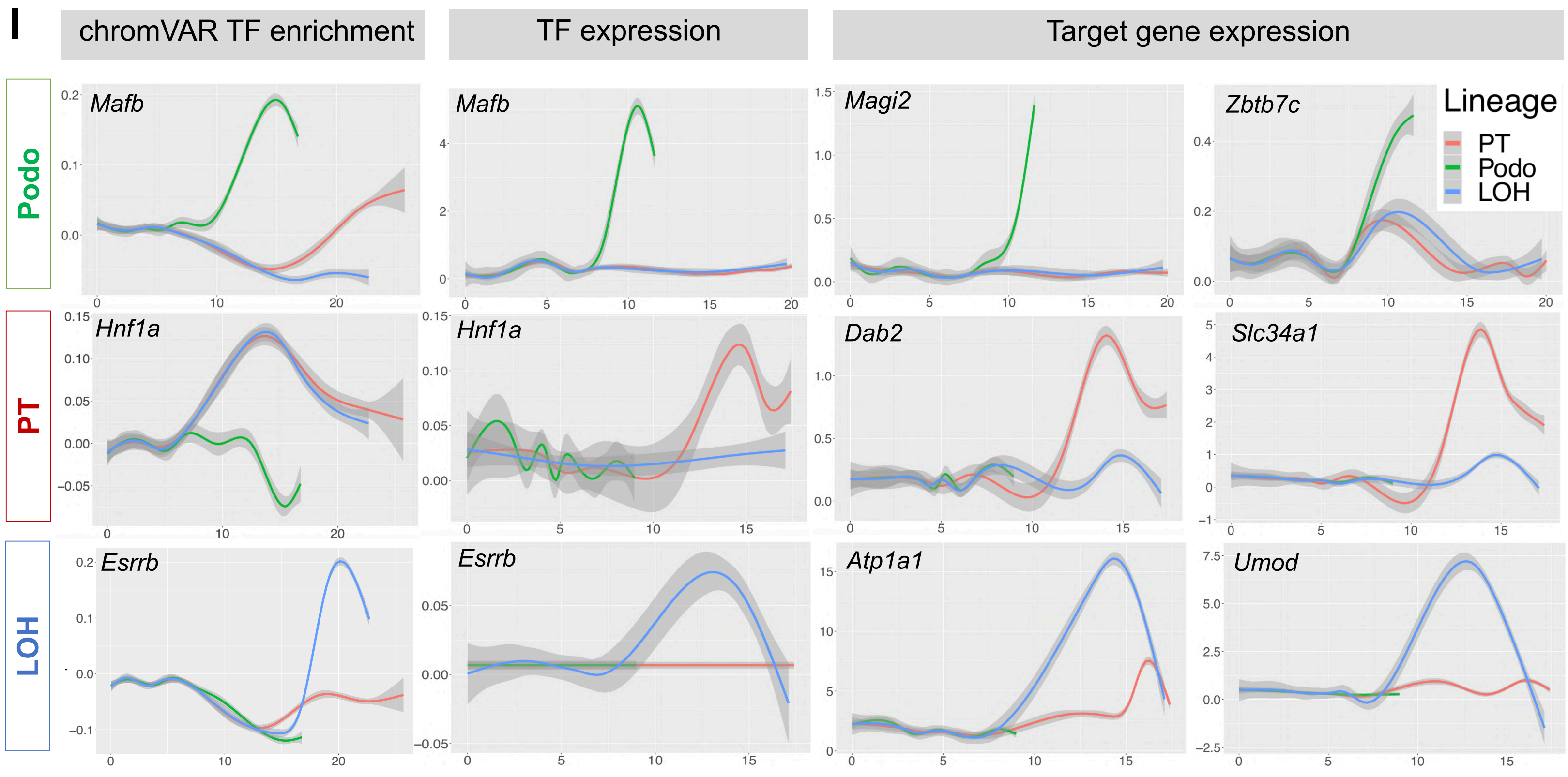
I
chromVAR TF enrichment
TF expression
Target gene expression
Podo
PT
LOH
Mafb
Magi2
Zbtb7c
Hnf1a
Dab2
Slc34a1
Esrrb
Atp1a1
Umod
Lineage
PT
Podo
LOH

#### Figure S4

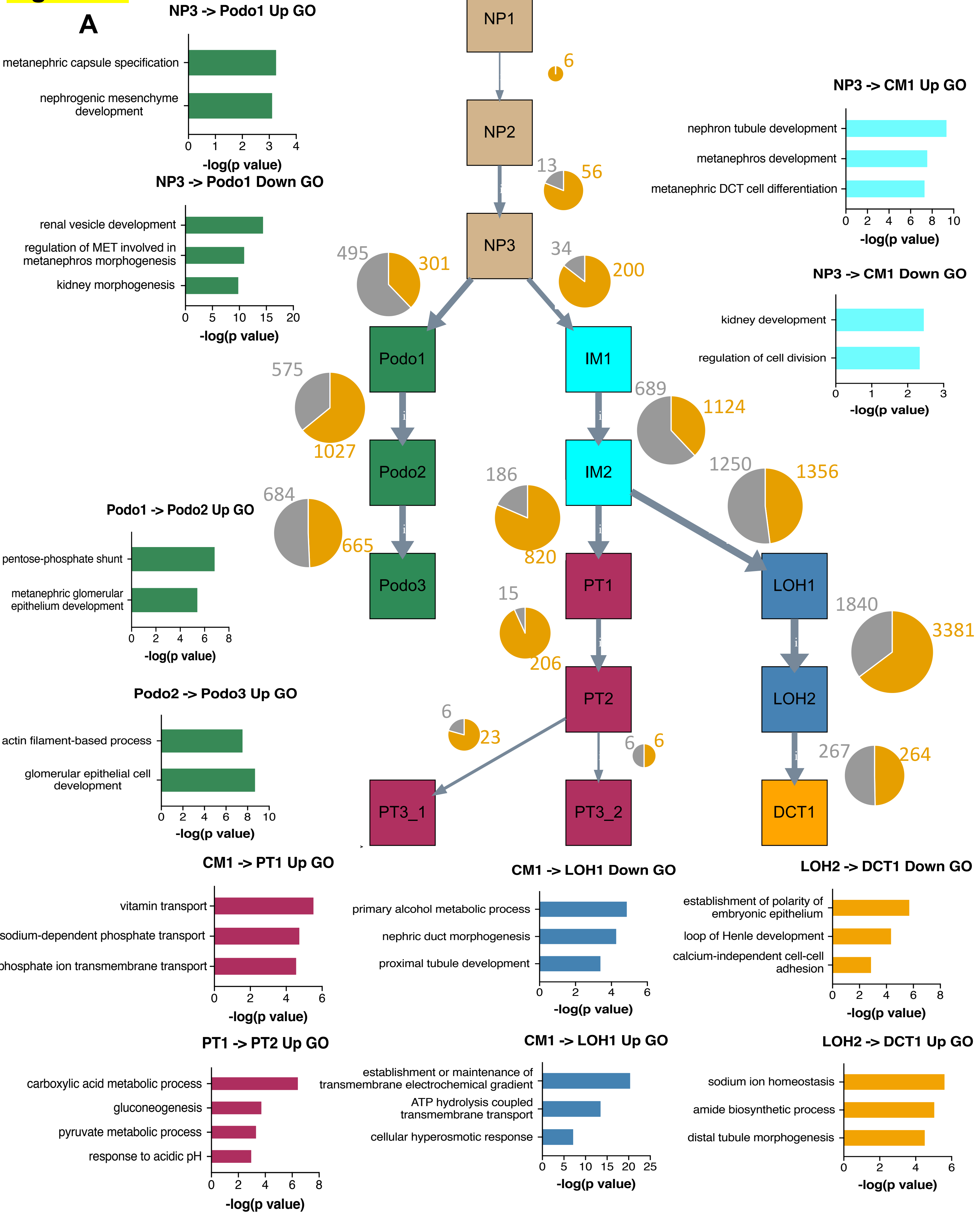
A
NP3 -> Podo1 Up GO
metanephric capsule specification
nephrogenic mesenchyme development
-log(p value)
NP3 -> Podo1 Down GO
renal vesicle development
regulation of MET involved in metanephros morphogenesis
kidney morphogenesis
-log(p value)
Podo1 -> Podo2 Up GO
pentose-phosphate shunt
metanephric glomerular epithelium development
-log(p value)
Podo2 -> Podo3 Up GO
actin filament-based process
glomerular epithelial cell development
-log(p value)
NP1
NP2
NP3
Podo1
Podo2
Podo3
IM1
IM2
PT1
PT2
PT3_1
PT3_2
LOH1
LOH2
DCT1
6
13
56
495
301
34
200
575
1027
689
1124
684
665
186
820
1250
1356
15
206
1840
3381
6
23
6
6
267
264
NP3 -> CM1 Up GO
nephron tubule development
metanephros development
metanephric DCT cell differentiation
-log(p value)
NP3 -> CM1 Down GO
kidney development
regulation of cell division
-log(p value)
CM1 -> PT1 Up GO
vitamin transport
sodium-dependent phosphate transport
phosphate ion transmembrane transport
-log(p value)
CM1 -> LOH1 Down GO
primary alcohol metabolic process
nephric duct morphogenesis
proximal tubule development
-log(p value)
LOH2 -> DCT1 Down GO
establishment of polarity of embryonic epithelium
loop of Henle development
calcium-independent cell-cell adhesion
-log(p value)
PT1 -> PT2 Up GO
carboxylic acid metabolic process
gluconeogenesis
pyruvate metabolic process
response to acidic pH
-log(p value)
CM1 -> LOH1 Up GO
establishment or maintenance of transmembrane electrochemical gradient
ATP hydrolysis coupled transmembrane transport
cellular hyperosmotic response
-log(p value)
LOH2 -> DCT1 Up GO
sodium ion homeostasis
amide biosynthetic process
distal tubule morphogenesis
-log(p value)

**Figure S4**

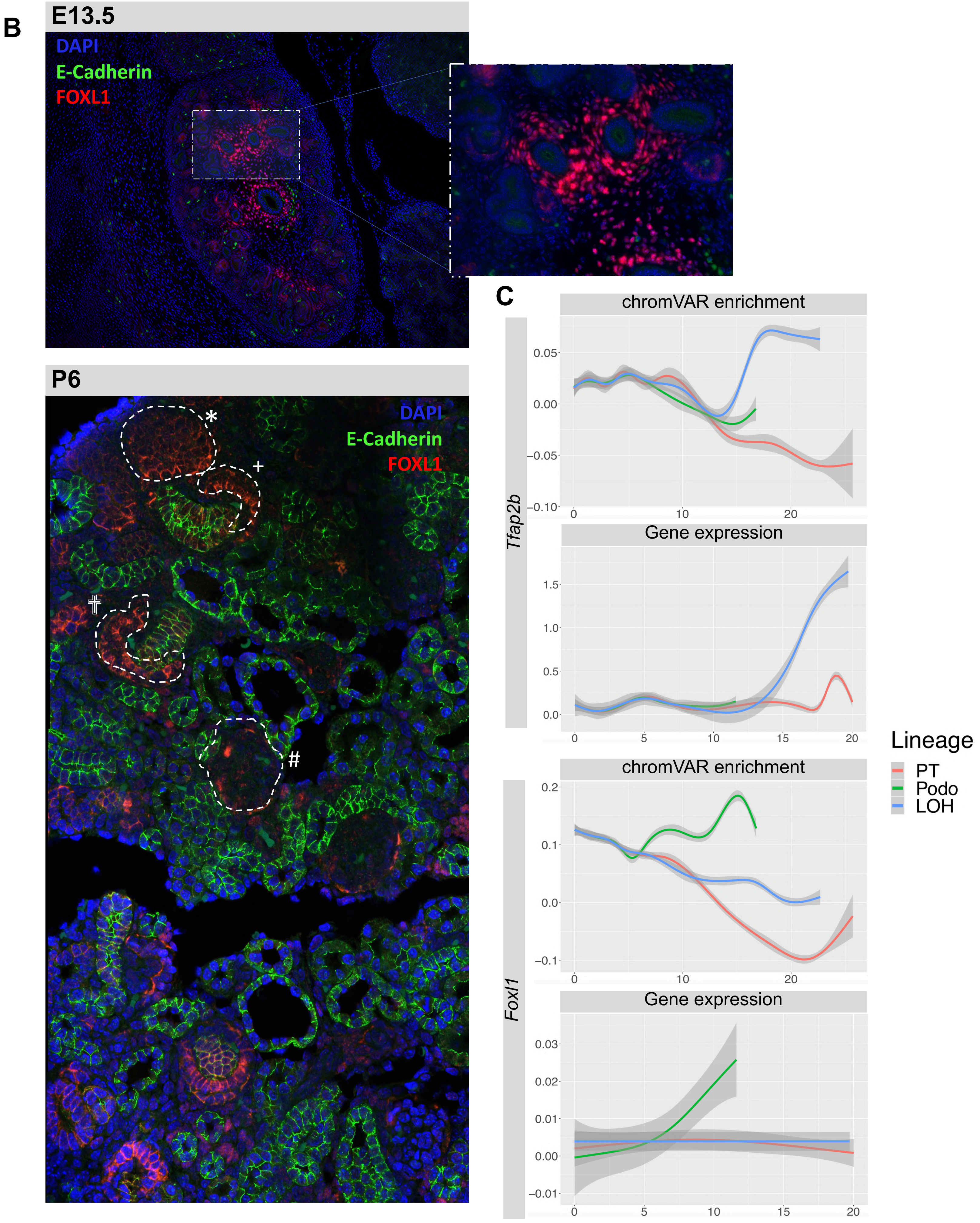

B
E13.5
DAPI
E-Cadherin
FOXL1
P6
DAPI
E-Cadherin
FOXL1
*
+
†
#
C
chromVAR enrichment
0.05
0.00
−0.05
−0.10
0
10
20
Tfap2b
Gene expression
1.5
1.0
0.5
0.0
0
5
10
15
20
chromVAR enrichment
0.2
0.1
0.0
−0.1
0
10
20
Foxl1
Gene expression
0.03
0.02
0.01
0.00
−0.01
0
5
10
15
20
Lineage
PT
Podo
LOH

**Figure S4**

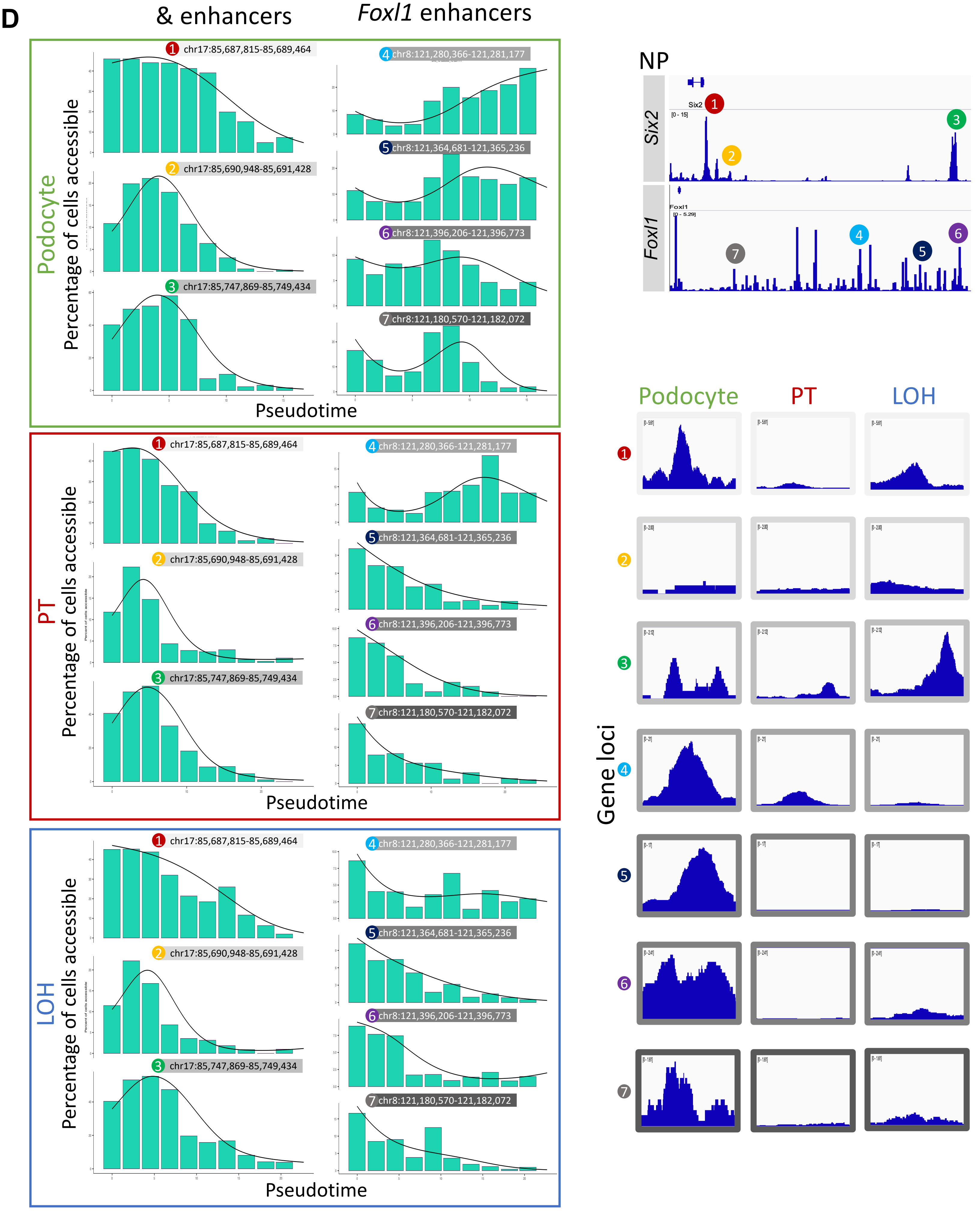
D
Six2 promoters & enhancers
Foxl1 enhancers
Podocyte
PT
LOH
Percentage of cells accessible
Pseudotime
1 chr17:85,687,815-85,689,464
2 chr17:85,690,948-85,691,428
3 chr17:85,747,869-85,749,434
4 chr8:121,280,366-121,281,177
5 chr8:121,364,681-121,365,236
6 chr8:121,396,206-121,396,773
7 chr8:121,180,570-121,182,072
NP
Six2
Foxl1
[0 - 15]
[0 - 5.29]
Gene loci

### Figure S5

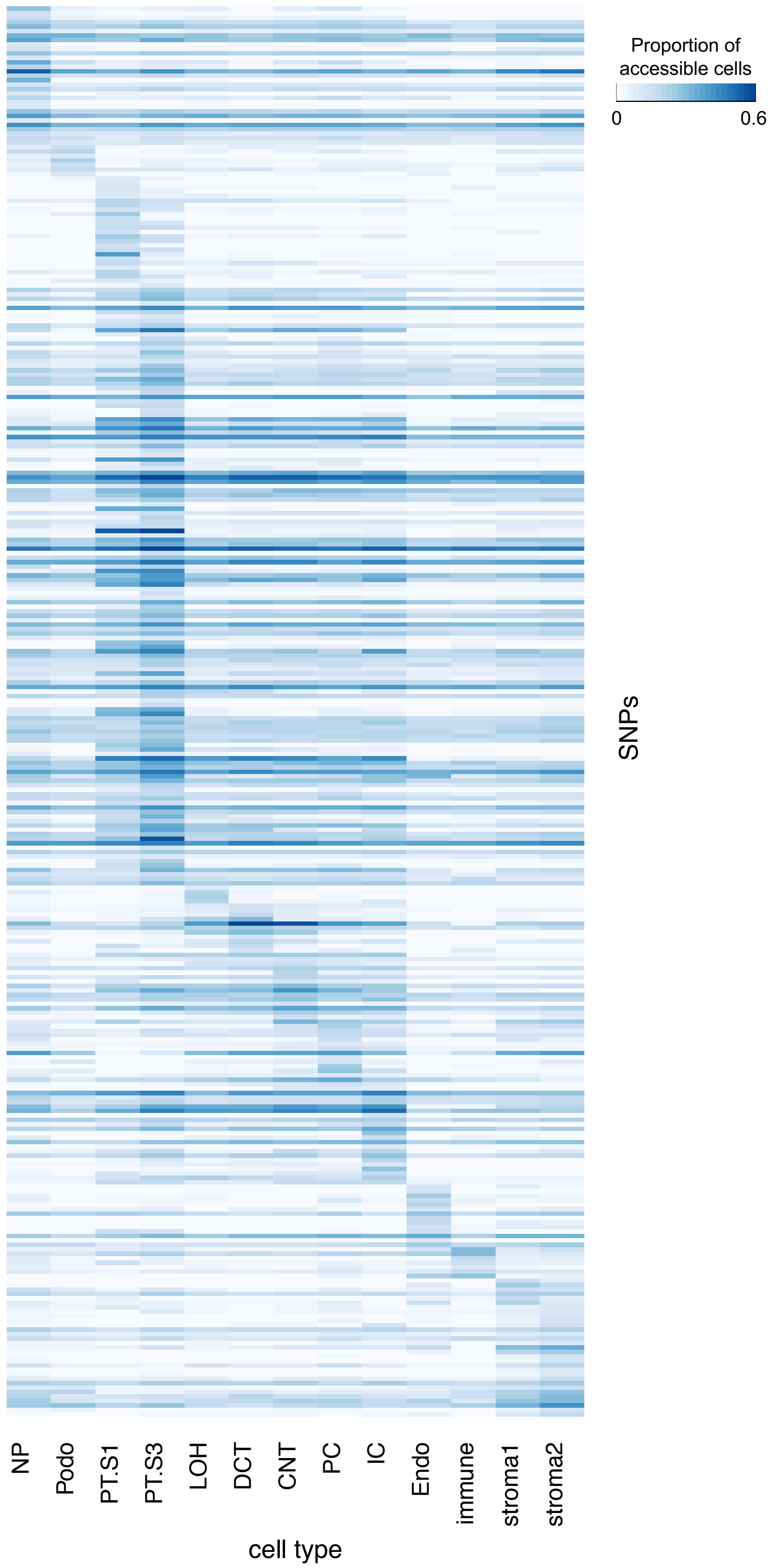

Proportion of accessible cells
0
0.6
SNPs
NP
Podo
PT.S1
PT.S3
LOH
DCT
CNT
PC
IC
Endo
immune
stroma1
stroma2
cell type

### Figure S6

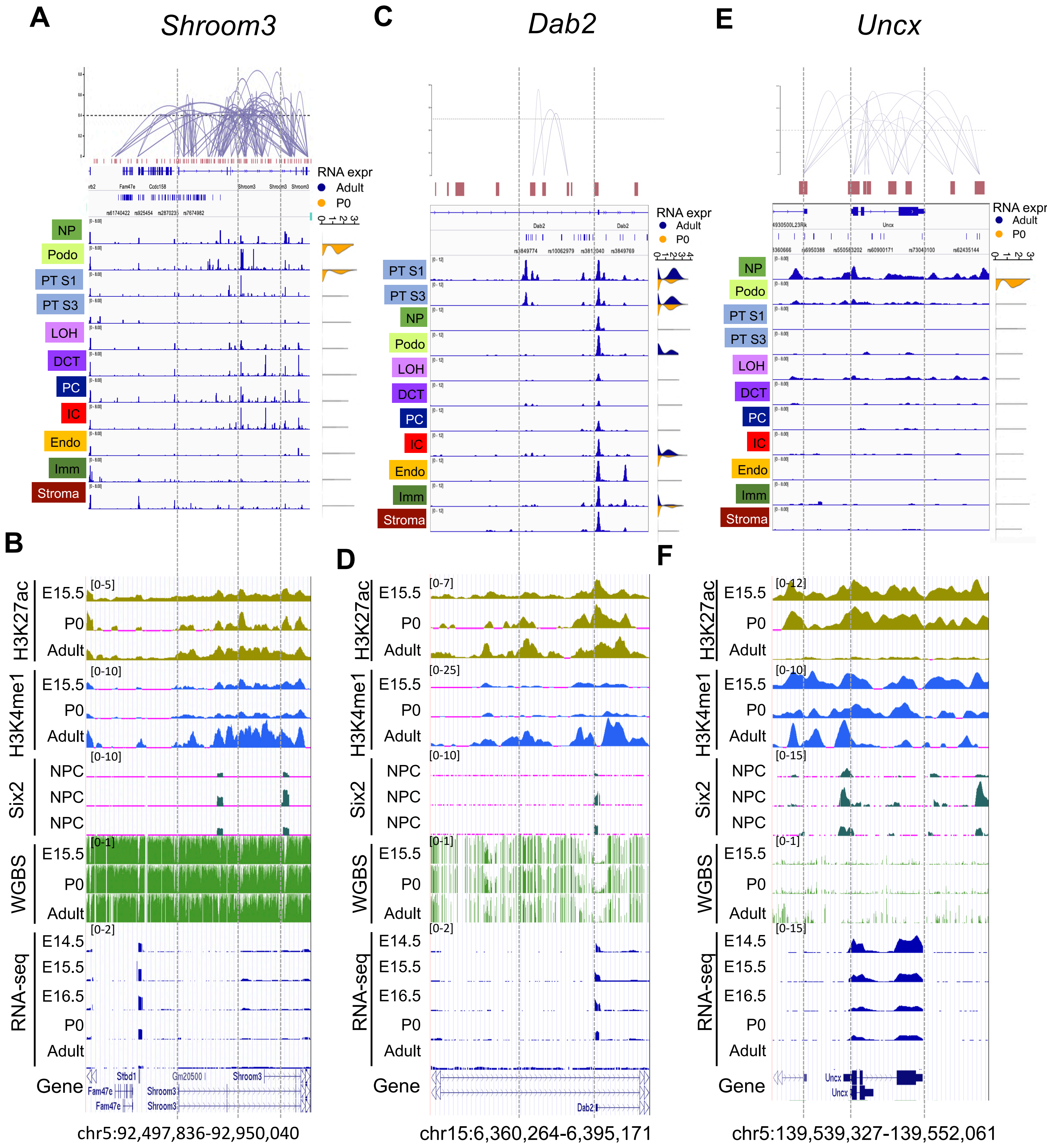

A
Shroom3
C
Dab2
E
Uncx
RNA expr
Adult
P0
NP
Podo
PT S1
PT S3
LOH
DCT
PC
IC
Endo
Imm
Stroma
B
D
F
H3K27ac
E15.5
P0
Adult
H3K4me1
Six2
NPC
WGBS
RNA-seq
E14.5
E16.5
Gene
chr5:92,497,836-92,950,040
chr15:6,360,264-6,395,171
chr5:139,539,327-139,552,061
